# Development of a Humanized Anti-Fibrotic Antibody Targeting Extracellular Collagen Assembly to Reduce Post-Traumatic Scarring

**DOI:** 10.64898/2026.04.20.719618

**Authors:** Andrew R. Mendelsohn, Bo Yu, Jolanta Fertala, James W. Larrick, Andrzej Fertala

## Abstract

**Background:** Excessive accumulation of fibrillar collagen causes pathological scarring and fibrosis. A promising anti-fibrotic strategy targets the extracellular assembly of collagen fibrils rather than intracellular synthesis pathways. We previously developed a chimeric monoclonal antibody targeting the C-terminal telopeptide of the α2(I) chain of human collagen I that effectively disrupts fibrillogenesis. This study details the engineering of a humanized antibody variant optimized for therapeutic application, augmented with a collagen-binding peptide (CBP) to enhance targeted retention in fibrotic tissues.

**Methods:** A humanized ACA was engineered by *in silico* homology modeling, complementarity-determining region grafting, and sequence optimization to eliminate chemical liabilities. Variants were expressed in mammalian cells and evaluated for binding kinetics and specificity. To improve spatial localization, the CBP was fused to the antibody. The lead variant was assessed for *in vitro* cytotoxicity, matrix retention, and in vivo efficacy using a rabbit model of post-traumatic knee arthrofibrosis.

**Results:** The humanized ACA variants maintained high specificity and affinity for the α2Ct target domain. Fusing the CBP to the C-terminus of the light chain (C-cbpACA) successfully enhanced matrix retention without compromising target engagement or causing cellular toxicity. In the rabbit arthrofibrosis model, intra-articular C-cbpACA delivery significantly reduced flexion contracture and decreased total collagen deposition in the joint capsule compared to untreated controls.

**Conclusion:** We successfully engineered a clinically viable, humanized, and matrix-targeted anti-fibrotic antibody that specifically inhibited extracellular collagen assembly and exhibited enhanced localization within fibrotic tissues. This construct represents a promising therapeutic strategy for mitigating pathological scarring and improving post-traumatic functional outcomes.

## INTRODUCTION

Scar formation is a fundamental biological process that repairs injured tissues by forming natural, collagen-based patches that preserve the structural integrity of damaged sites. In a balanced healing response, scar tissue undergoes progressive remodeling that does not significantly impair normal physiological function. In contrast, excessive scarring leads to abnormal fibrotic deposits that disrupt tissue mechanics and compromise organ function (1).

Inflammation drives both adaptive and pathological scarring by promoting the release of potent pro-fibrotic factors. These factors activate fibroblastic cells, stimulating the synthesis and deposition of extracellular matrix components, particularly fibrillar collagens. During normal healing, inflammation and fibroblast activation resolve over time; however, in fibrotic conditions, collagen deposition persists, resulting in progressive tissue stiffening and dysfunction.

Our research group focuses on post-traumatic fibrotic scarring of musculoskeletal and neuronal tissues, including joint capsules, skeletal muscle, and peripheral nerves. Severe scarring of the joint capsule results in arthrofibrosis, a condition characterized by progressive joint stiffness and profound restriction of mobility (2, 3). Similarly, fibrotic remodeling of injured peripheral nerves can impair axonal regeneration, leading to persistent neuromuscular dysfunction (4, 5).

Across diverse anatomical and etiological contexts, the formation of collagen-rich deposits drives excessive scarring. Collagen I constitutes the dominant structural component of scar tissue, accounting for approximately 80% of its dry mass (6). Consequently, many anti-fibrotic therapeutic strategies aim to reduce collagen I production or accumulation. Canonical approaches typically target intracellular signaling pathways that regulate collagen biosynthesis. However, despite extensive investigation, the development of safe and effective anti-fibrotic therapeutics remains challenging (7).

To expand the repertoire of viable anti-fibrotic targets, we focus on the extracellular assembly of collagen fibrils, a process essential for scar maturation. Collagen fibrillogenesis occurs through the aggregation of individual collagen I molecules, driven by a highly specific interaction between the telopeptide domains of one collagen molecule and the telopeptide-binding region (TBR) of another. This interaction initiates fibril formation, producing a robust, enzymatically cross-linked scaffold characteristic of mature scar tissue (8, 9).

We have previously demonstrated that this critical interaction can be disrupted by targeting the telopeptide domains of collagen I. Specifically, the C-terminal telopeptide of the pro-α2(I) chain (α2Ct) represents an attractive molecular target for limiting collagen fibrillogenesis. Building on this concept, we developed a monoclonal anti-α2Ct antibody (ACA) and validated its safety and efficacy in reducing fibrotic scarring across multiple experimental models (2, 10–12).

In the present study, we engineered and characterized a humanized ACA optimized for therapeutic application. We outline the critical steps involved in antibody humanization and developability optimization, and further enhance spatial localization and functional performance by fusing the antibody with a collagen-binding peptide (CBP). Together, the structural and functional data presented here establish a robust foundation for evaluating this rationally engineered, matrix-targeted ACA variant in clinically relevant models of fibrotic disease.

## MATERIAL AND METHODS

### Engineering anti-collagen antibody (ACA) variants

#### Anti-fibrotic target

Our group pioneered an anti-fibrotic strategy that specifically targets extracellular collagen fibrillogenesis. We demonstrated that inhibiting collagen fibril assembly by disrupting the interaction between the α2Ct and TBR effectively limits scar formation (2, 8, 9, 11). The following sections describe the key steps involved in developing a clinically relevant antibody-based inhibitor of fibrotic scarring (see summary in Supplementary Materials, Fig. S1).

#### Prototypic ACA

Using cDNA sequences encoding the original mouse-derived ACA’s heavy and light chains, the recombinant prototypic ACA was generated as a chimeric IgG_1_ variant. The chimeric ACA (referred to as ch1ACA) consisted of mouse-derived variable regions and human-derived constant regions corresponding to the heavy γ1 chain and the light κ chain (8, 11–13). *In vitro* and biological models confirmed the target specificity and the potential of ch1ACA to limit collagen fibrillogenesis (12, 14). Complete cDNA and amino acid sequences of the chimeric chains are available in our patent document (15).

### Humanization of ACA

#### *In silico* design and antibody sequence optimization

To generate clinically viable anti-collagen variants targeting the α2Ct domain (Fig. 1), a 3D homology model of the murine variable regions was constructed to guide the humanization and framework engineering strategy.

**Figure 1.**
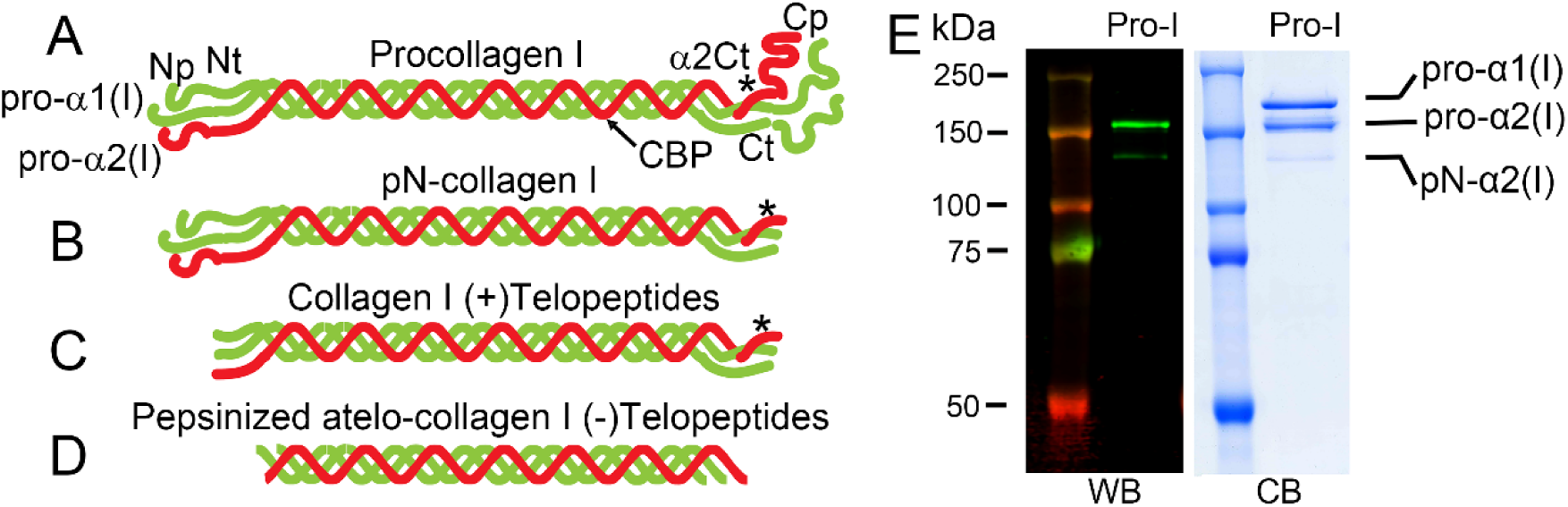
Schematic of collagen I variants and demonstration of ACA binding specificity. (A) Intact procollagen I molecule comprising pro-α1(I) and pro-α2(I) chains, complete with N- and C-terminal propeptides (Np, Cp) and telopeptides (Nt, Ct). An asterisk () denotes the pro-α2(I) C-terminal telopeptide (α2Ct), which contains the epitope recognized by the ACA antibody. The collagen-binding peptide (CBP) recognition site is also indicated. (B) Partially processed pN-collagen I, which lacks the C-terminal propeptide but retains the N-terminal propeptide and telopeptide regions (*). (C) Processed collagen I lacking both propeptides but retaining the telopeptide domains (*). (D) Pepsin-digested collagen I lacking all propeptides and telopeptides. (E) Representative Western blot (WB) demonstrating ACA specificity for the α2Ct target in intact pro-α2(I) and partially processed pN-α2(I) chains. ACA does not bind the pro-α1(I) chain, which lacks the α2Ct epitope. A Coomassie blue (CB)-stained gel of procollagen I chains (Pro-I) is included for reference.

The prototypic murine heavy chain, derived from the mouse germline VH9-2, was humanized by grafting its complementarity-determining regions (CDRs) onto two distinct human acceptor frameworks: VH1-2 and VH7-4-1. This strategy generated six distinct heavy-chain variants (H1-H6).

Similarly, the prototypic murine light chain, derived from the mouse germline VK8-21, was grafted onto the human VK4-1 acceptor framework, generating two light chain variants (L1 and L2). These specific human acceptor frameworks were selected for their high sequence and structural homology with the original murine scaffolds.

Because direct humanization often reduces target affinity, the 3D model was used to identify structurally critical murine framework residues. Specific back mutations to these murine residues were designed and incorporated into the human frameworks to preserve the precise architecture of the antigen-binding sites. Concurrently, the engineered sequences were evaluated to identify and eliminate inherent chemical liabilities that could compromise long-term developability.

#### Heavy chain optimization

The murine CDRs were grafted onto two distinct human framework scaffolds to evaluate optimal structural support. Variants H1, H2, and H3 were engineered using the human VH7-4-1 framework, while variants H4, H5, and H6 were engineered using the human VH1-2 framework.

Developability screening identified the asparagine-threonine (NT) motif in CDR2, a known deamidation-prone (16). To mitigate this risk, variants H3 (from the VH7-4-1 lineage) and H6 (from the VH1-2 lineage) were selected as foundational scaffolds. Targeted amino acid substitutions were introduced to mutate the asparagine (N) to either a serine (S) or an alanine (A) within CDR2.

This engineering strategy successfully removed the NT motif, with the H3 scaffold yielding optimized variants H7 and H8, and the H6 scaffold yielding optimized variants H9 and H10.

#### Light chain optimization

For light-chain humanization, murine CDRs were grafted onto the human VK-4-1 framework scaffold to generate variants L1 and L2. Sequence analysis identified a high-risk asparagine-serine (NS) motif within CDR1. This motif poses a significant risk of non-enzymatic deamidation, leading to product heterogeneity and a subsequent loss of target affinity. To mitigate the risk of functional degradation during manufacturing and storage, variant L2 was selected as the foundation for further engineering. A targeted amino acid substitution was introduced to replace asparagine (N) with alanine (A) in CDR1, successfully removing the NS motif and generating the optimized variant L3.

Ultimately, these combined structural and developability engineering strategies yielded a robust panel of lead ACA variants designed to balance high target engagement with the chemical stability required for therapeutic advancement.

#### Molecular cloning and expression

To evaluate the functional characteristics of these designs, a series of mammalian expression vectors was constructed. First, a chimeric anti-collagen antibody was generated to establish a baseline for target binding and activity (Fig. S1).

By expressing the wild-type murine variable domains linked to human constant regions, this chimeric construct confirms that the selected mammalian expression system and the human IgG4 isotype do not inherently disrupt antigen recognition.

Consequently, the modified chimeric ACA version, referred to as ch4ACA, served as the definitive reference point; any subsequent loss of binding affinity observed in the fully humanized variants can be specifically attributed to the framework grafting and CDR modifications rather than the assay conditions or constant domain class switching.

#### Chimeric heavy chain

The wild-type murine ACA VH sequence was cloned into the LB601 expression vector, which encodes a human γ4 constant region.

#### Chimeric light chain

The wild-type murine ACA VK sequence was cloned into the LB603 vector to express the chimeric human κ light chain.

Following the establishment of the chimeric baseline, the humanized constructs were cloned into the corresponding expression vectors:

#### Humanized heavy chains

The synthesized humanized ACA VH genes (variants H1 through H6) were cloned into the LB602 vector for expression as human heavy γ4 chains.

#### Humanized light chains

The humanized ACA VK constructs (variants L1 and L2) were cloned into the LB604 vector for expression as human κ light chains.

#### Generation of stable production cell lines

Following the initial screening (see below), the lead humanized ACA variants, H8L2 and H9L2, were advanced for stable cell line generation (Fig. S1). To support continuous, high-yield manufacturing, stable production cells for these variants were generated using a CHO-S host cell line (Thermo Fisher Scientific, Waltham, MA).

#### The SwiMR expression system

To facilitate rapid development and isolation of highly productive clones, the cDNAs encoding the H8L2 and H9L2 variants were inserted into a proprietary SwiMR expression system (**17**). This system utilizes a switchable membrane reporter to enable the efficient isolation of high-producing cells via fluorescence-activated cell sorting (FACS; Figs. S2 & S3).

Using this expression system, the antibody heavy and light chains are expressed from two separate vectors, each driven by a strong, constitutive human EF1α promoter to ensure robust expression in mammalian cells.

#### Heavy chain vector (LB602)

The heavy chain sequences (H8 and H9), complete with signal peptides, were expressed using the LB602 vector (Fig. S2). This vector encodes an engineered human IgG_4_ Fc region with specific mutations (S228P/F234A/L235A) that enhance stability while minimizing effector functions. Crucially, LB602 incorporates an internal ribosome entry site (IRES)-mediated bicistronic expression cassette downstream of the heavy chain, which encodes a membrane-anchored green fluorescent protein (GFP). Because the heavy chain and GFP are co-transcribed, GFP fluorescence intensity serves as a direct, quantifiable reporter of antibody expression level. To accommodate downstream clinical development, this IRES-GFP cassette is flanked by LoxP sites, allowing for its targeted removal from the host chromosome via treatment with recombinant Cre DNA recombinase. The LB602 vector utilizes a puromycin resistance gene for mammalian selection.

#### Light chain vector (LB604)

The light chain sequence (L2) and its signal peptide were expressed using the LB604 SwiMR expression vector. For dual-selection compatibility, this vector carries a neomycin (G418) resistance gene (Fig. S2).

#### Selection of stable transfectants

Before transfection, the LB602 and LB604 expression plasmids were linearized with ScaI. Then, the linearized vectors were co-transfected into CHO-S cells maintained in serum-free CD FortiCHO media using a lipid-based transfection reagent (Thermo Fisher Scientific).

To establish stable integrants, the transfected pools were subjected to rigorous dual-antibiotic selection with 10 µg/ml puromycin and 500 µg/ml G418 for 2 weeks. Following the selection phase, FACS isolated the top ∼1% of cells with the highest GFP reporter signals, yielding sorted pools of approximately 200,000 highly productive cells. These enriched pools, designated as 44.42.2bp1 for H8L2 and 44.42.3bp1 for H9L2, were subsequently expanded to support preclinical characterization *in vitro* and *in vivo* (Fig. S3).

#### Engineering ACA variants with a collagen-binding peptide (CBP)

To enhance retention of ACA at collagen-rich injury sites, antibody variants incorporating a collagen-binding peptide (TKKTLRT) were engineered (**18**). The CBP was fused at one of three locations on the H9L2 scaffold: the C-terminus of the light chain (C-cbpACA), upstream of the hinge (U-cbpACA), or downstream of the hinge (D-cbpACA), using flexible (G_4_S) linkers (Fig. 2). For the C terminus of the light chain the CBP was linked with (G_4_S)_2_. CBP was linked upstream of the hinge with one G_4_S linker. CBP was linked downstream of the hinge flanked by G_4_S linkers on both sides.

**Figure 2.**
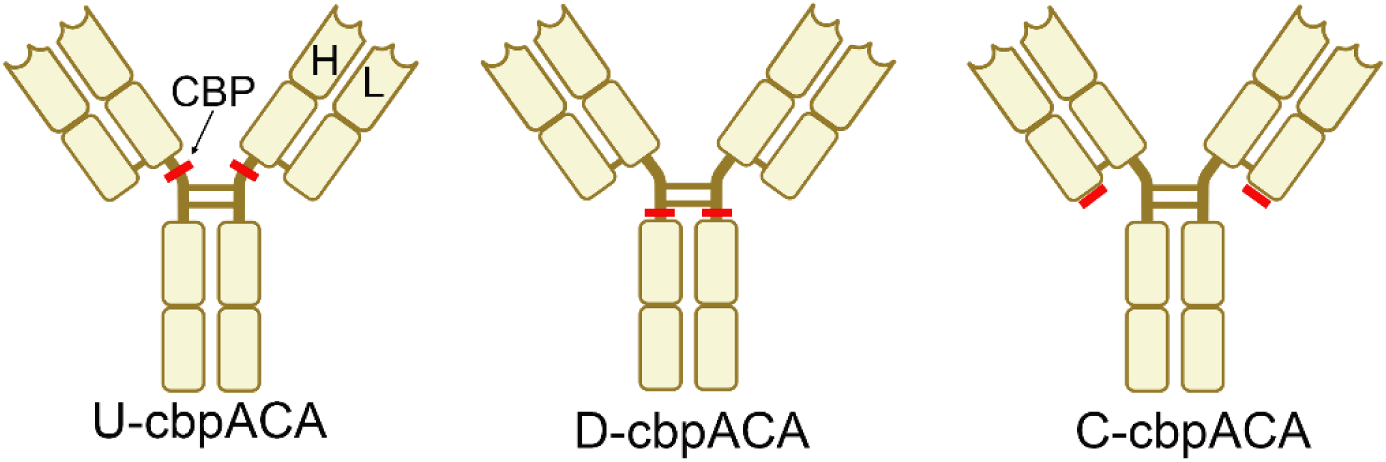
Schematic of collagen-binding peptide (CBP) placement in ACA constructs. The diagram illustrates CBP positioned upstream (U-cbpACA) or downstream (D-cbpACA) of the heavy (H) chain hinge region, or at the C-terminus (C-cbpACA) of the light (L) chain.

### Biochemical and functional characterization of the ACA variants

As indicated above, we followed rational steps to engineer humanized ACA variants, which included (Fig. S1):

1. The IgG1 version of a prototypic chimeric ACA construct (ch1ACA)
2. The IgG4 variant of chimeric ACA (ch4ACA)
3. A set of 12 humanized first-generation ACA constructs that included H1-H6 heavy chains and L1-L2 light chains (h1ACA)
4. Modified humanized second-generation ACA constructs that included H7-H10 heavy chains and a combination of L2-L3 light chains (h2ACA)
5. H9L2-based third-generation constructs with CBP cassette (cbpACA)

Thus, we tested the above constructs to determine their binding kinetics, specificity, and the ability to block collagen fibrillogenesis. These tests allowed us to select an optimal candidate to act as an anti-fibrotic agent in a relevant animal model of arthrofibrosis.

#### Antibody integrity and homogeneity

We used electrophoresis to assess the integrity and chain assembly of the ACA variants. Moreover, we evaluated their homogeneity using size-exclusion chromatography (SEC). Specifically, we used the BioBasic SEC-300 high-performance liquid chromatography (HPLC) column (Thermo Fisher Scientific) equilibrated with phosphate-buffered saline (PBS) at a flow rate of 1 ml/min. Protein elution profiles were monitored at 280 nm.

#### Apparent binding affinities of the h1ACA variants

To determine the apparent binding affinity (defined as the antibody concentration required to saturate 50% of target sites, or EC_50_), we utilized an enzyme-linked immunosorbent assay (ELISA). First, plates were coated with streptavidin (2 µg/ml) and blocked to prevent non-specific binding. Then, a synthetic biotinylated α2Ct target peptide (0.1 µg/ml) was captured on the plates for 1 hour. After removing the unbound peptide, h1ACA variants were added in 2-fold serial dilutions, starting at 2 µg/ml. Peptide-bound variants were detected using a horseradish peroxidase (HRP)-conjugated anti-human Fc antibody. Absorbance values were plotted against the antibody concentration to determine the EC_50_.

#### Biosensor-based assays of the h1ACA-α2Ct binding kinetics

Binding kinetics for the ch4ACA (control) and 8 h1ACA variants selected from the initial 12 candidates based on EC_50_ values of approximately 0.1 µg/ml were analyzed using an optical biosensor (Octet, Sartorius, Göttingen, Germany). Curia (Hayward, CA) performed these assays.

First, streptavidin-coated biosensors (Octet SA Biosensors, Sartorius) were loaded with biotinylated α2Ct target peptide. Next, association rates (k_on_) were measured by exposing sensors to the ACA variants at concentrations ranging from 4.7 nM to 300 nM. For each ACA concentration, the dissociation rates (k_off_) were measured after exposing the sensor to ligand-free solvent. Equilibrium dissociation constants (K_D_) were calculated using a monovalent binding model (K_D_ = k_off_/k_on_).

#### ACA-procollagen I binding kinetics

While the synthetic α2Ct target peptide enabled an initial assessment of the h1ACA constructs’ binding affinities, the next essential step for selecting variants with optimal binding properties was characterizing the binding to the native target, procollagen I.

In brief, procollagen I was purified from rabbit dermal fibroblast cultures, biotinylated, and immobilized on streptavidin-coated sensors (Pioneer SensiQ, Sartorius) (19, 20). Binding kinetics were assessed by exposing the sensors to 12 h1ACA variants at concentrations ranging from 6.25 nM to 200 nM. Moreover, ch1ACA and ch4ACA constructs served as controls.

#### Binding specificity

To confirm the specificity of 12 h1ACA variants for binding the α2Ct target in the context of the intact α2(I) chain, we employed a multichannel Western blot approach that allows simultaneous analysis of up to 24 antibodies.

In brief, denatured procollagen I was loaded into a single well spanning the width of a 7.5% polyacrylamide gel, while a molecular mass marker was loaded into a separate standard well. After electrophoresis, the separated pro-α1(I) and pro-α2(I) chains were transferred onto a nitrocellulose membrane. Then, the membrane was blocked and clamped into an MPX Blotting System (LI-COR Biosciences, Lincoln, NE, USA). Next, individual antibody variants were added to separate channels and incubated with the membrane for 2 hours. After washing the channels, antibody-procollagen complexes were visualized using goat anti-human IgG antibodies conjugated with a green chromophore (IRDye 800CW, LI-COR Biosciences).

As a negative control, we assessed binding of the ACA variants against pepsin-digested collagen I, which lacks the intact α2Ct epitope found in native procollagen I molecules (Fig. 1). Similar assays were performed using the prototypic ch1ACA and ch4ACA variants.

#### Binding kinetics of the h2ACA constructs

The binding kinetics of the h2ACA variants to native procollagen I were measured using a biosensor, as described above.

#### Testing the H8L2 and H9L2 constructs

As described above, we selected H8L2 and H9L2 constructs as lead candidates for production and *in vivo* tests. The following points describe detailed assays of these constructs:

#### Inhibition of collagen fibrillogenesis *in vitro*

The H8L2 and H9L2 antibodies were evaluated for their potential to inhibit de novo collagen fibril formation. We applied the in vitro methods previously described by Chung et al. (8). Briefly, purified procollagen I was converted to collagen I using LysC enzyme (1 µg/ml) (Sigma-Aldrich, St. Louis, MO) at 25°C for 30 minutes. Under these conditions, LysC digests procollagen propeptides while preserving crucial collagen telopeptides, including the antibody’s α2Ct epitope. At this stage, collagen molecules remain in a non-aggregated, (i.e., non-fibrillar) form. Subsequently, LysC was inhibited with Nα-p-tosyl-L-lysine chloromethyl ketone (TPCK, Sigma-Aldrich) and added to a final concentration of 1 mM.

The H8L2 and H9L2 antibodies or control non-specific human IgG (hIgG, Sigma-Aldrich) were added to collagen samples at five concentrations ranging from 3 nM to 270 nM. Given the collagen concentration of 240 nM, the molar collagen-to-antibody ratios in the samples ranged from approximately 1:1 to 1:0.01.

Collagen fibril formation was initiated by incubating the samples at 37°C for 24 hours. The samples were then centrifuged for 30 minutes to separate large fibrillar aggregates, which settle in the pellet (P), from non-aggregated monomers remaining in the supernatant (S). Subsequently, the P and S fractions were analyzed by electrophoresis in a 10% polyacrylamide gel under denaturing and reducing conditions. Proteins were visualized by staining with Coomassie blue, and collagen bands were quantified by densitometry (Odyssey CLx, LI-COR Biosciences).

Subsequently, the P-to-S signal ratios were calculated for each sample and plotted against antibody concentration; high ratios indicated that most collagen molecules were in the fibrillar P fraction, whereas low ratios indicated that collagen molecules remained primarily as non-aggregated monomers in the S fraction. Results of these assays were expressed as the mean ± SD of three technical replicates.

#### Microscopic assays of collagen fibrils

To complement the electrophoretic analysis of fibril formation, samples were visualized using dark-field (DF) microscopy (Eclipse E600, Nikon Inc., Melville, NY) (10). Following incubation at 37°C for 24 hours, aliquots of the collagen-antibody mixtures were transferred to glass slides and observed under dark-field illumination. Under these conditions, collagen fibrils appear as distinct, needle-shaped structures. Conversely, the absence of visible structures indicates that the aggregation process was effectively blocked. Notably, DF microscopy detects only large aggregates (fibrils); non-aggregated collagen monomers remain in the solution and are not visible.

### Analyses of the cbpACA constructs

#### cbpACA variants’ binding kinetics

The binding kinetics of the cbpACA variants were measured using a biosensor, as described above. As indicated, these variants included the α2Ct-binding sites within their CDRs, as well as the binding site for the triple-helical region of the collagen α2(I) chain provided by the CBP cassette.

To study specific contributions of these two sites to interactions with their intended targets, namely the triple helical domain of collagen I (for CBP) and the α2Ct domain (for antibody CDRs), we prepared sensors coated with the following targets:

1. Pro-I: native procollagen I (Fig. 1), produced by cultured rabbit dermal fibroblasts. This target included procollagen N-terminal and C-terminal propeptides (NP, CP), telopeptides (Nt, Ct), and the triple-helical domain (TH). We expected Pro-I to bind the cbpACA variants via the α2Ct and TH domains.
2. Pro-I-peps: native procollagen I digested with pepsin (atelo-collagen I). Pepsin digests globular propeptides and part of telopeptides (Fig. 1); TH remains intact. We expected this target to bind cbpACA variants via α2Ct (if incompletely digested) and TH.
3. Tendon collagen I: collagen (predominantly collagen I), which we extracted from rabbit tendon using pepsin. We predicted this target to bind cbpACA variants via α2Ct (if incompletely digested) and TH.
4. Mouse collagen I: Collagen (predominantly collagen I) isolated by us from the mouse skin using acetic acid extraction (with no pepsin). Our previous research (unpublished) demonstrated that mouse α2Ct has poor binding affinity for our antibody. Hence, we expected this collagen to bind tested antibodies predominantly via TH-CBP interaction.
5. Chick collagen II: a commercial collagen II isolated from the chick sternum (Sigma-Aldrich). We predicted that this collagen type would bind the antibodies predominantly via interaction with the TH-CBP cassette, because of relatively high homology between the α2(I) chain MMP1 cleavage site region (the CBP target) and the corresponding region of the α1(II) chain.
6. Synthetic α2Ct: a human sequence-based ACA target. We expected this target to bind via the α2Ct-binding sites of the cbpACA variants.

To perform binding studies, the above targets were biotinylated, then immobilized on streptavidin-coated sensors.

After determining the concentrations of the C-cbpACA, U-cbpACA, and D-cbpACA variants, we prepared their serial dilutions ranging from 6.25 to 400 nM. Subsequently, the binding kinetics for the above targets were measured for each antibody construct. Finally, the k_on_ and k_off_ rates were calculated and used to determine the K_D_ values for each cbpACA variant-target pair.

Additionally, the ACA constructs lacking the CBP domain served as controls.

#### Measurement of C-cbpACA retention in cell culture conditions

Based on the binding kinetics, we selected the C-cbpACA variant for analysis of its key characteristics before proceeding to pilot animal studies (see Results section). First, we analyzed the behavior of this antibody within the collagen-rich matrix produced by rabbit dermal fibroblasts.

The fibroblasts were seeded on 8-well slides or 24-well plates at a density of 5 × 10^4^ cells. Then, to develop a collagen-rich matrix, the cells were cultured for 3 days in Dulbecco’s Modified Eagle’s Medium (DMEM) supplemented with 10% fetal bovine serum (FBS) and 40 µg/ml of L-ascorbic acid phosphate magnesium salt n-hydrate (WAKO Inc.).

After that time, we confirmed the presence of the matrix produced by cells seeded onto 8-well slides by fixing the cell layers with methanol and staining the collagenous matrix using polyclonal anti-collagen antibodies (AFCol-1) and the secondary anti-rabbit IgG antibodies conjugated to Alexa Fluor 594 (Thermo Fisher Scientific) (12).

To measure the retention of unmodified H9L2 ACA and C-cbpACA, these antibodies were added to established cell cultures at 200 µg/ml. Cell culture medium was changed every 2 days, and cultures with antibodies were maintained for 8 days.

Following this incubation period, the cell monolayers were washed extensively to remove unbound antibodies. Next, the cell layers were lysed with a lysis buffer to assess the presence of antibodies trapped within collagen-rich matrices. Subsequently, proteins present in the lysates were separated by electrophoresis. We analyzed the presence of the antibodies by Western blot using anti-human IgG primary antibodies conjugated with a green chromophore (IRDye 800CW, LI-COR Biosciences). Pixel intensities of the antibody-specific heavy (Hγ) and light (Lκ) chains were measured by densitometry. Finally, the results of these assays were plotted using data from 3 technical repeats.

We also measured glyceraldehyde 3-phosphate dehydrogenase (GAPDH) as an internal protein-loading control using a mouse anti-GAPDH antibody (Santa Cruz Biotechnology, Dallas, TX) and a goat anti-mouse secondary antibody conjugated with a red chromophore (IRDye 680RD, LI-COR Biosciences).

#### Evaluation of cytotoxicity of the C-cbpACA variant

We used the CytoSelect™ MTT Cell Proliferation kit (Cell Biolabs Inc., San Diego, CA) to assess the potential toxicity of the C-cbpACA variant. This assay relies on the ability of the MTT reagent (3-(4,5-dimethylthiazol-2-yl)-2,5-diphenyl-2H-tetrazolium bromide) to pass through cellular and mitochondrial membranes of living cells. Once inside, the reagent is reduced to formazan, a violet-blue-colored molecule. Cell viability is then determined by measuring the formazan-dependent absorbance at 570 nm.

#### Assay of acute cytotoxicity

We evaluated the acute cytotoxicity of the C-cbpACA variant using rabbit dermal fibroblasts (RF) and human keloid-derived fibroblasts (HF) as follows:

- RF and HF cells were seeded in triplicate into 96-well plates at a density of 1 × 10^4^ cells per well.
- The cells were cultured in DMEM supplemented with 10% FBS.
- To assess acute toxicity, the cells were treated for 24 hours with C-cbpACA at concentrations up to 300 μg/ml.
- Following treatment, the amount of solubilized formazan was measured at 570 nm with a plate reader (Benchmark Plate Reader, Bio-Rad Laboratories, Hercules, CA).
- The means were plotted from 3 technical repeats (±SD) using GraphPad Prism software (GraphPad Software Inc., Boston, MA).

#### Long-term cytotoxicity assay

To investigate the long-term effect of C-cbpACA on cell proliferation, RF and HF cells were seeded at 3 × 10³ and 1.5 × 10³ cells/ml, respectively. The cells were subsequently cultured for 7 days in DMEM containing 10% FBS and 200 μg/ml C-cbpACA. Proliferation was measured colorimetrically using the MTT assay at designated time points over 7 days.

Two control groups were used:

- negative control, where cells were cultured without the antibody.
- positive control, where cells were cultured with 0.5 μg/ml of actinomycin-D, an anti-proliferative agent, to confirm toxicity effects.

The means from 3 technical repeats (±SD) were plotted as a percentage of the absorbance measured on day one.

### Animal studies

#### Rabbit arthrofibrosis model

The Thomas Jefferson University Institutional Animal Care and Use Committee (IACUC) approved the study protocol (protocol #02108-2).

Given the pilot nature of this in vivo study with the C-cbpACA construct, we utilized five New Zealand White (NZW) rabbits (three males, two females; aged 8 to 12 months; Charles River, Wilmington, MA) to analyze the antibody’s impact on key parameters of collagen-rich deposits in a post-traumatic arthrofibrosis model (2, 14, 21). Because we previously reported the outcomes for untreated controls (CTR) and rabbits treated with ACA variants lacking the CBP cassette, these groups were omitted from the analysis (2, 14). Yet, we utilized the results of these historical studies to assess the impact of the C-cbpACA studied here. This experimental design minimized redundancy and reduced the total number of animals required for this preliminary investigation.

Our rabbit model of arthrofibrosis meets essential criteria for a valid preclinical animal model. The rabbit musculoskeletal system shares fundamental features with that of humans, including the composition and structure of bones, ligaments, tendons, and muscles. Additionally, rabbit and human collagenous proteins exhibit similar patterns of biosynthesis and fibrillogenesis. Importantly, the therapeutic target in this study, the α2Ct, is identical between rabbits and humans. This model has been successfully applied by numerous independent research groups, including our own (2, 6, 14, 21–23).

#### Blood collection and analysis

Blood samples were collected at baseline and 4, 8, and 12 weeks after the initial surgery. Complete blood counts (CBCs) were performed using the Element HT5 instrument (Antech Diagnostics, Inc., Levittown, PA).

#### Surgical procedures

Post-traumatic contracture was generated by simulating an intra-articular fracture and bleeding (using drill holes) and a posterior capsule (PC) injury.

The contralateral uninjured limb served as the control. A refillable pump (see below) was installed subcutaneously to deliver C-cbpACA directly into the articular cavity of the injured knee joints. In brief, an incision in the middle of the right knee was made starting 5 mm proximally to the patella and extending distally to the tibial tubercle and down the tibia. Subsequently, a lateral parapatellar arthrotomy was made, and the patella was dislocated medially. After flexing the knee, the fat pad was released and retracted medially. Next, the anterior and posterior cruciate ligaments (ACL and PCL) were transected, and the knee was hyperextended to 45° to disrupt the PC.

We drilled a 2-mm drill hole in the medial femoral condyle and a second 2-mm hole beginning as far posteriorly in the notch as possible and exiting through the lateral condyle. Starting from the notch, a custom-made silicone catheter (0.6 mm ID/1.2 mm OD, Access Technologies, Skokie, IL) with a fixed retention bead at one end was threaded through the hole. While the retention bead secured the catheter inside the operated knee, the opposite end was connected to a pump.

To exacerbate the injury, the injured knee was maintained in flexion with a Kirschner wire (K-wire) for 8 weeks. In brief, a drill was driven in the anterior-posterior direction to create a 2-mm bicortical drill hole in the tibia. Subsequently, a 1.6-mm K-wire (Modern Grinding, Port Washington, WI) was passed through the hole in the tibia (Fig. 3). Flexing the knee, the K-wire was maneuvered to exit through the femur’s lateral side incision. Two 90° bends, approximately 1 cm apart, were made to form a hook at the end of the K-wire. Next, the hook was positioned around the femur. Finally, the knee was fixed at flexion by tightening a nut on the opposite, threaded end of the K-wire.

**Figure 3.**
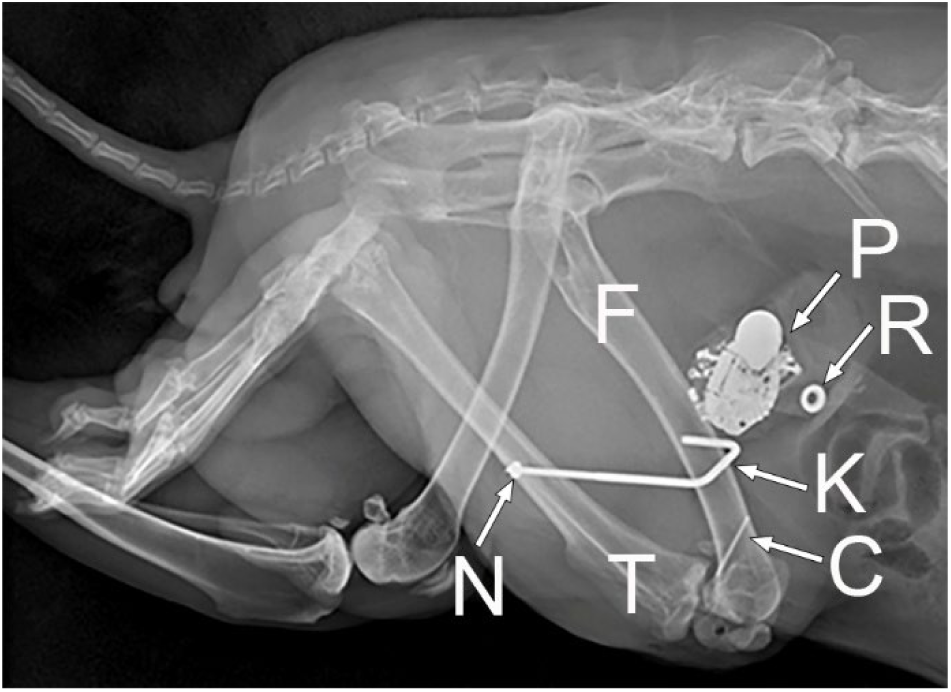
Radiograph of a rabbit following surgical induction of arthrofibrosis. Anatomical landmarks and surgical hardware are indicated as follows: T, tibia; F, femur; P, pump; Kw, Kirschner wire; Tc, tube connector; Rp, refill port.

#### Pump installation

Using a subcutaneous, programmable pump (the iPrecio SMP-200, Primetech Corp., Tokyo, Japan) allowed us to control delivery of the antibody directly to the knee injury site. The pump was placed into a subdermal pocket created during surgery. Using a stainless-steel tube connector (Access Technologies, Skokie, IL), the free end of the catheter was connected to a silicon tube attached directly to the refillable pump. The pump was filled with 0.9 ml of the antibody solution at 30 mg/ml. Nylon sutures (Ethicon Inc., Somerville, NJ) were tied around the silicon tubes at their juncture to ensure the stability of the connection site. Finally, the incisions were closed with 4-0 absorbable sutures (Redisorb, MYCO Medical, Cary, NC). Following surgery, the rabbits were allowed unrestricted cage activity.

#### K-wire removal

After 8 weeks of immobilization, the rabbits underwent a second surgery to remove the K-wires. The heterotopic ossification along the K-wire was manually disrupted. Subsequently, the rabbits were allowed 4 weeks of unrestricted cage time, after which they were euthanized.

#### Antibody delivery

In addition to the initial portion of the antibody supplied by the pump implanted at the initial surgery, 4 weeks later, the pump was refilled with an additional portion (0.9 ml) of C-cbpACA at a concentration of 30 mg/ml.

#### Histology of tissues and organs

At euthanasia, various organs and tissues (e.g., brain, heart, liver, tendons, nerves) were collected, processed for histology, and stained with hematoxylin and eosin (H&E) to assess structural and inflammatory changes possibly due to treatment with C-cbpACA (2).

#### Mechanical assays of the flexion contracture

Joint immobilization and knee injury lead to stiffness and flexion contracture, primarily due to PC scarring. Thus, measuring contracture provides a net clinically relevant endpoint to assess C-cbpACA efficacy in modulating collagen fibrillogenesis (6, 14).

Flexion contracture was measured using a custom-made instrument (Test Resources, Inc., Shakopee, MN; Fig. S4) (2, 14, 24). In brief, the injured or the contralateral uninjured leg was fixed in the instrument with the tibia and femur positioned at a right angle. This position was considered the starting point of zero degrees. Subsequently, applying the 40°/min loading rate, an extension torque of 0.2 Nm was applied. Upon reaching the value of 0.2 Nm, the angle corresponding to the joint extension was recorded. The injured knee flexion contracture score was then expressed as the uninjured control knee extension angle divided by the injured contralateral knee extension angle (2, 6, 14, 24). Hence, relatively low ratio values indicated relatively small flexion contracture, while large ratio values indicated more severe flexion contracture.

#### Collagenous PC matrix

Since PC scarring drives knee stiffness in our arthrofibrosis model, the PCs were dissected for histological, biochemical, and spectroscopic assays focused on collagenous material (2, 6, 14, 25).

#### Histological assays of collagen fibrils

Histological sections of the PCs were stained with H&E to visualize tissue morphology and cellularity. Additionally, similar sections were stained with picrosirius red, a collagen-specific dye. By combining this staining method with polarized-light microscopy, we could approximate the fibrils’ thickness, organization, and packing density (6, 26–28). As the thickness and packing of fibers increase, their birefringence color changes from green to yellow to orange to red (i.e., from shorter to longer wavelengths) (26, 29–32).

Employing a polarizing microscope (Eclipse LV100POL, Nikon Inc.) and the NIS Elements software (Nikon Inc.), the following groups of the birefringence colors were defined in captured images: (i) green birefringence (GB, identifies thin, loosely packed fibrils), (ii) yellow birefringence (YB, identifies intermediate-thickness fibrils), and (iii) red birefringence (RB, identifies thick, tightly packed fibrils) (2, 6, 14).

All fibrils in captured viewing areas were analyzed (6, 33). In brief, the above birefringence colors were defined by applying the software’s “color threshold” function. Subsequently, the software determined the areas occupied by pixels corresponding to the defined colors. Considering the sum of all fibers to be 100%, the percentages for each color group in the analyzed samples were calculated. The exact threshold settings were applied to all images to ensure consistent definition of birefringence colors. Automated calculations of all fibrils in the viewing areas eliminated any potential bias.

#### Fourier transform infrared spectroscopy (FTIR)

FTIR spectroscopy of collagen-rich tissues provides information about their protein and proteoglycan composition (34). Thus, we used this method to measure the relative collagen content in the scar tissue formed in injured PCs of antibody-treated and CTR rabbits. As a non-collagenous reference, we selected a peak corresponding to sulfated glycosaminoglycans (GAGs), a common non-collagenous component of collagen-rich connective tissues. The collagen-GAG ratios were calculated based on the areas of the spectral peaks corresponding to collagen, centered around 1338 cm^-1^ wavenumber (ν) and the sulphated GAGs, centered around 1064 cm^-1^ ν (2, 35, 36).

Since the maturity of collagen cross-links often increases during the fibrotic process, its assays provide a relevant parameter to evaluate the fibrotic status of collagen-rich deposits formed in the presence of the C-cbpACA (37–39). Consequently, we analyzed the maturity of collagen cross-links by measuring their mature pyridinoline (PYR) form and immature dehydro-dihydroxynorleucine (de-DHLNL) form. The maturity of collagen fibrils was expressed as the PYR-de-DHLNL ratio. The trivalent PYR cross-link peak centers around 1660 cm^-1^ ν, and the immature divalent deDHLNL cross-link peak centers around 1690 cm^-1^ ν (2, 40, 41).

To perform FTIR assays, paraffin-embedded 5-µm-thick tissue sections were deposited on the MirrIR low-e microscope slides (Kevley Technologies, Chesterland, OH). Subsequently, an FTIR spectrometer was used to analyze regions of interest (ROIs) corresponding to the PC collagen-rich areas (Spotlight 400, PerkinElmer, Waltham, MA). The measurements were done in the 4000 cm^-1^ to 748 cm^-1^ ν spectral range, at a pixel resolution of 50 μm, 8 scans per pixel, and a spectral resolution of 4 cm^-1^ ν. The Spectrum Image software generated co-added spectra from scanned ROIs (PerkinElmer, Inc.).

In all assays of the FTIR-derived spectra, overlapping peaks were deconvoluted and analyzed based on the second-order derivative spectra and pre-determined bell-type Gaussian peak fitting function using the OriginLab software (version 2025, OriginLab Corporation, Northampton, MA, USA) (2, 42, 43).

#### Total collagen content

To further analyze the impact of C-cbpACA treatment on collagen content in scar tissue formed in PCs, we quantified collagen-specific hydroxyproline (HP) in the isolated PCs. In brief, PC samples were frozen in liquid nitrogen and then pulverized in a stainless-steel mortar. Next, the samples were treated with a 3:1 mixture of chloroform and methanol to extract lipids and then lyophilized and weighed. Then, tissues were hydrolyzed in 6 N HCl. Subsequently, we measured the HP content using an HP kit (Sigma-Aldrich, St Louis, MO) (6, 44). Finally, the HP content was used to calculate the collagen content per unit of tissue dry mass.

#### Statistical analysis

Analysis was conducted to evaluate differences in analyzed parameters across the three rabbit cohorts: (i) historical untreated CTR, (ii) historical standard ACA-treated group, and (iii) the C-cbpACA pilot group created here (2, 14, 45, 46).

Due to the highly unbalanced study design, specifically the exceptionally small C-cbpACA pilot group (n = 4) compared to the larger historical CTR and ACA cohorts, violations of normality and homogeneity of variance required a nonparametric Independent-Samples Kruskal-Wallis H test to compare the distributions of continuous scores across the three categorical groups.

Post-hoc comparisons were adjusted using Bonferroni correction. Statistical significance was defined as p < 0.05.

## RESULTS

### First-generation h1ACA constructs demonstrate proper structure and target binding properties

Following transient expression of 12 h1ACA variants (H1-H6 with L1 and H1-H6 with L2), we analyzed their purity, structural integrity, and target binding properties.

#### Electrophoretic assays

Electrophoretic analysis of the ACA constructs under reducing conditions demonstrated the proper migration patterns of the heavy and light chains. Moreover, the presence of a single protein band in the non-reducing condition indicated proper stabilization of the ACA variants’ heavy and light chains via disulfide bonds (Fig. S5).

#### SEC HPLC analyses

All h1ACA variants eluted as a single peak with retention times corresponding to a predicted molecular mass of 150 kDa (Fig. S6).

#### Apparent binding affinity

Utilizing the α2Ct peptide as a target, ELISA demonstrated concentration-dependent binding of all h1ACA variants (Fig. S7). Based on the binding results, we selected 8 constructs with an EC_50_ of ∼0.1 µg/ml. Those constructs included: H2L1, H3L1, H4L1, and H6L1, as well as H2L2, H3L2, H5L2, and H6L2. Subsequently, the binding kinetics of these constructs were analyzed using a biosensor. In those assays, we used a synthetic α2Ct as a binding target. Table I presents the results.

**Table. I.** Binding kinetics of the ACA variants with EC_50_ ∼0.1 µg/ml (α2Ct used as binding target)

| ACA variant | $k_{on}$ [$M^{-1}s^{-1}$] | $k_{off}$ [$s^{-1}$] | $K_D$ [M] |
| --- | --- | --- | --- |
| ch4ACA | 2.90E+04 | <1.0E-07 | <1.0E-12 |
| H2/L1 | 3.52E+04 | 1.36E-05 | 3.86E-10 |
| H2/L2 | 5.94E+04 | 6.59E-05 | 1.11E-09 |
| H3/L1 | 1.85E+04 | <1.0E-07 | <1.0E-12 |
| H3/L2 | 5.67E+04 | <1.0E-07 | <1.0E-12 |
| H4/L1 | 4.15E+04 | 3.64E-05 | 8.77E-10 |
| H5/L2 | 6.43E+04 | 5.86E-05 | 9.12E-10 |
| H6/L1 | 6.27E+04 | 2.45E-06 | 3.91E-11 |
| H6/L2 | 7.14E+04 | 1.10E-05 | 1.54E-10 |

#### Assays of binding kinetics using native procollagen I

To further characterize the binding kinetics of the h1ACA variants, we used procollagen I as a native target. Table II presents the results of the biosensor-based binding assays.

**Table. II.** Binding kinetics of the ACA variants (procollagen I used as binding target)

| ACA variant | $k_{on}$ [$M^{-1}s^{-1}$] | $k_{off}$ [$s^{-1}$] | $K_D$ [M] |
| --- | --- | --- | --- |
| H1/L1 | 7.86E+03 | 6.55E-04 | 8.34E-08 |
| H2/L1 | 4.94E+03 | 7.94E-04 | 1.61E-07 |
| H3/L1 | 1.33E+04 | 1.71E-04 | 1.28E-08 |
| H4/L1 | 1.14E+04 | 6.64E-04 | 5.85E-08 |
| H5/L1 | 1.04E+04 | 2.06E-04 | 1.98E-08 |
| H6/L1 | 1.63E+04 | 2.01E-04 | 1.23E-08 |
| H1/L2 | 9.73E+03 | 1.93E-04 | 1.99E-08 |
| H2/L2 | 1.47E+04 | 6.30E-04 | 4.27E-08 |
| H3/L2 | 1.20E+04 | 1.04E-04 | 8.66E-09 |
| H4/L2 | 7.82E+03 | 2.66E-04 | 3.40E-08 |
| H5/L2 | 1.16E+04 | 2.72E-04 | 2.33E-08 |
| H6/L2 | 1.59E+04 | 1.11E-04 | 6.96E-09 |
| ch4ACA | 1.63E+04 | 1.39E-04 | 8.51E-09 |
| ch1ACA | 1.48E+04 | 1.04E-04 | 6.97E-09 |

#### Analysis of the binding specificity of the h1ACA antibodies

To ensure the h1ACA variants bound specifically to their intended target (i.e., α2Ct present in the context of the native pro-α2(I) chain), we used Western blot assays. As indicated in Fig. 4, all variants correctly recognized the α2Ct within pro-α2(I) chain as their target. Consistent with our earlier observations, the h1ACA variants also recognized a partially degraded pro-α2(I) chain form, pN-α2(I), in which the C propeptide was processed but the α2Ct epitope remained intact (8). In contrast, none of the variants recognized pepsin-digested procollagen I, in which propeptides and telopeptides were degraded. As expected, similar results were obtained with ch4ACA control (Fig 4).

**Figure 4.**
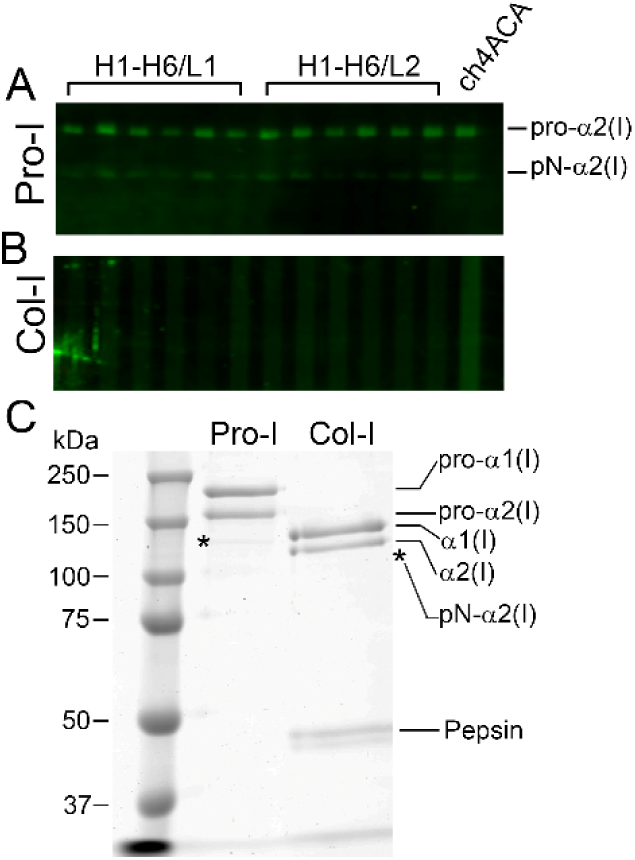
Binding specificity of first-generation h1ACA variants. Western blot analysis evaluating the interaction of h1ACA variants (H1-H6 heavy chains paired with L1 or L2 light chains) and the ch4ACA variant with (A) intact procollagen I chains and (B) pepsinized collagen I chains lacking the ACA-specific epitope. (C) Reference Coomassie blue-stained gel showing intact procollagen I (Pro-I) and pepsinized collagen I (atelo-collagen I). Intact chains (pro-α1(I), pro-α2(I)), mature chains (α1(I), α2(I)), and the partially processed pN-α2(I) chain (*) are indicated. Bands corresponding to the pepsin used for digestion are also marked.

### Second-generation h2ACA constructs show correct structure, integrity, high-affinity binding properties, and effectively inhibit collagen fibrillogenesis

#### Electrophoretic and chromatographic analyses

Purified h2ACA variants were first analyzed using electrophoretic and chromatographic methods. Results showed the expected electrophoretic migration of these variants (Fig. 5). Their chromatographic behavior was similar to that of the h1ACA counterparts (Fig. S6).

**Figure 5.**
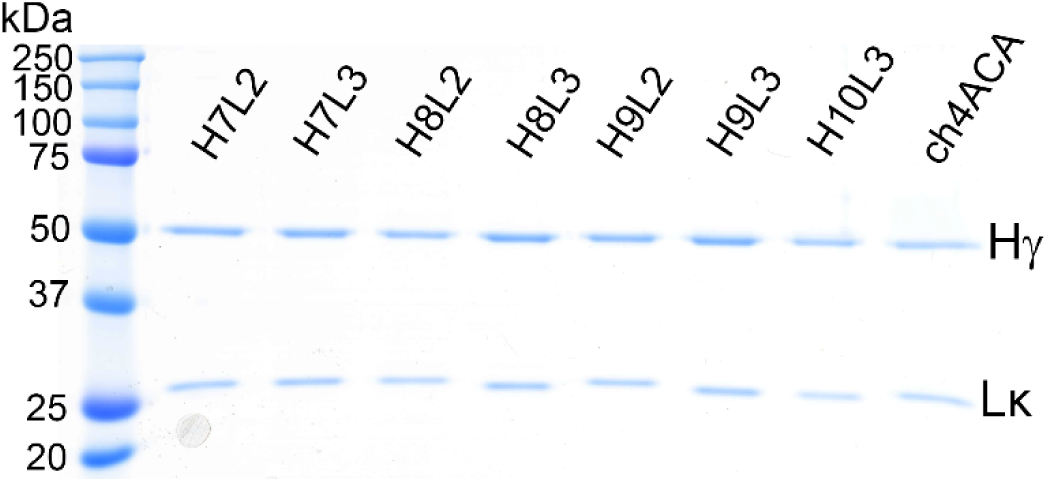
Electrophoretic analysis of second-generation h2ACA variants and the ch4ACA construct. The expected migration of the heavy (Hγ) and light (Lκ) chains confirms the correct structural assembly of these antibodies.

#### Binding kinetics of the h2ACA constructs

Biosensor-based assays revealed that all four h2ACA constructs retained high-affinity binding to native procollagen I (Table III). Based on favorable kinetic profiles, including low dissociation constants and balanced association/dissociation rates, we selected the H8L2 and H9L2 variants as lead candidates for functional and in vivo studies. As ACA variants with the L3 chain showed poor binding properties (Table III), we excluded them from further tests.

**Table. III.**
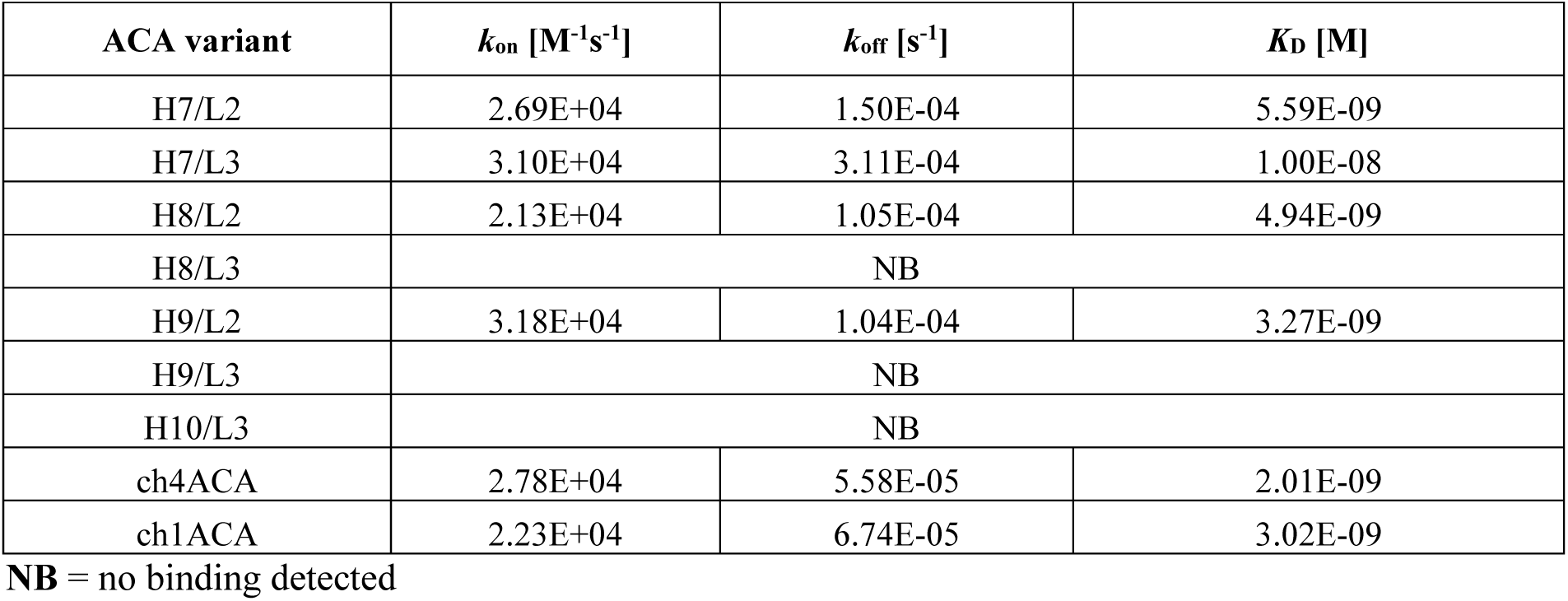
Binding kinetics of the ACA variants (procollagen I used as binding target)

| ACA variant | $k_{on}$ [$M^{-1}s^{-1}$] | $k_{off}$ [$s^{-1}$] | $K_D$ [M] |
| --- | --- | --- | --- |
| H7/L2 | 2.69E+04 | 1.50E-04 | 5.59E-09 |
| H7/L3 | 3.10E+04 | 3.11E-04 | 1.00E-08 |
| H8/L2 | 2.13E+04 | 1.05E-04 | 4.94E-09 |
| H8/L3 | NB |  |  |
| H9/L2 | 3.18E+04 | 1.04E-04 | 3.27E-09 |
| H9/L3 | NB |  |  |
| H10/L3 | NB |  |  |
| ch4ACA | 2.78E+04 | 5.58E-05 | 2.01E-09 |
| ch1ACA | 2.23E+04 | 6.74E-05 | 3.02E-09 |

**NB** = no binding detected

#### Inhibition of collagen fibrillogenesis in vitro

The H8L2 and H9L2 variants were evaluated for their ability to inhibit in vitro collagen fibrillogenesis. Electrophoretic analysis confirmed their inhibitory potential (Fig. 6).

**Figure 6.**
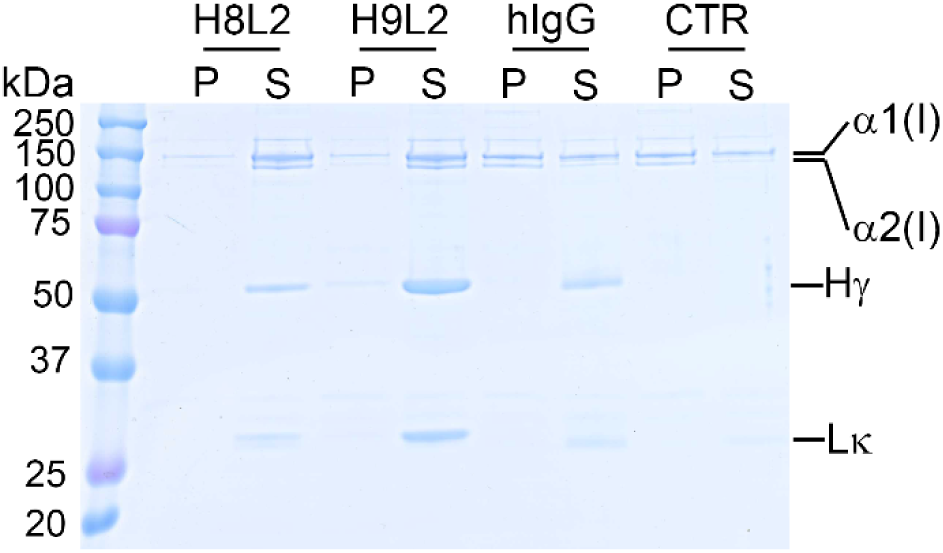
Electrophoretic analysis of collagen fibril formation in the presence of inhibitory antibodies H8L2 and H9L2. Controls include a non-inhibitory antibody (hIgG) and a no-antibody sample (CTR). Following fibril formation, samples were separated by centrifugation into fibrillar pellet (P) and non-aggregated monomeric supernatant (S) fractions prior to electrophoresis. In the presence of inhibitory antibodies, collagen I remains predominantly in the non-fibrillar monomeric form (S). In contrast, a robust fibrillar fraction (P) is observed with the non-inhibitory hIgG and CTR. Residual monomers (S) present in the uninhibited controls reflect the critical concentration threshold of collagen; fibril assembly ceases once the concentration of free collagen molecules falls below this limit. Collagen I α1(I), and α2(I) chains, and ACA’s heavy (Hγ) and light (Lκ) chains are indicated.

Moreover, we demonstrated a concentration-dependent inhibition by both variants, as evidenced by a progressive reduction in collagen content in the fibril-derived pellet fraction at higher antibody concentrations. In contrast, the control antibody (hIgG) failed to alter fibril assembly under identical conditions, regardless of concentration (Figs. 7 and 8).

**Figure 7.**
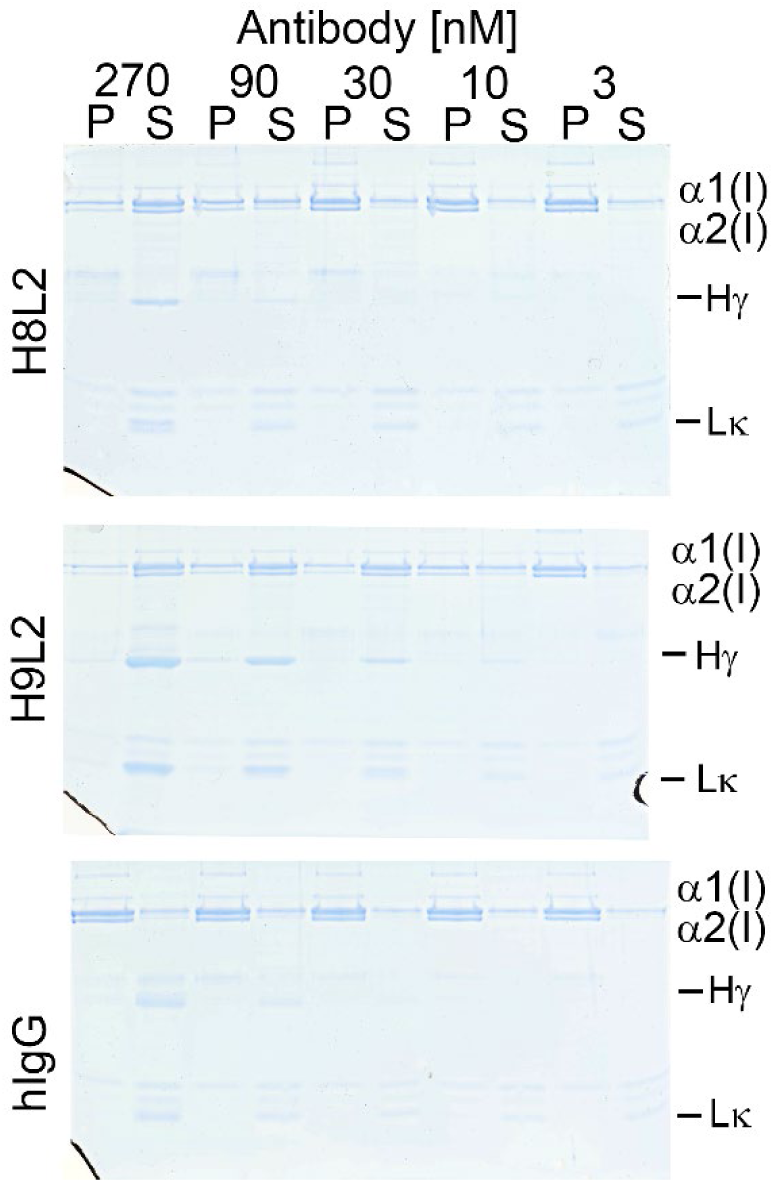
Electrophoretic analysis of collagen fibril formation in the presence of inhibitory antibodies H8L2 and H9L2 or a non-inhibitory antibody (hIgG). The fibrils formed at different antibody concentrations. Following fibril formation, samples were separated by centrifugation into fibrillar pellet (P) and non-aggregated monomeric supernatant (S) fractions prior to electrophoresis. In the presence of inhibitory antibodies, collagen I remains predominantly in the non-fibrillar monomeric form (S). In contrast, a robust fibrillar fraction (P) is observed with the non-inhibitory hIgG and CTR. Residual monomers (S) present in the uninhibited controls reflect the critical concentration threshold of collagen; fibril assembly ceases once the concentration of free collagen molecules falls below this limit. Collagen I α1(I), and α2(I) chains, and ACA’s heavy (Hγ) and light (Lκ) chains are indicated.

**Figure 8.**
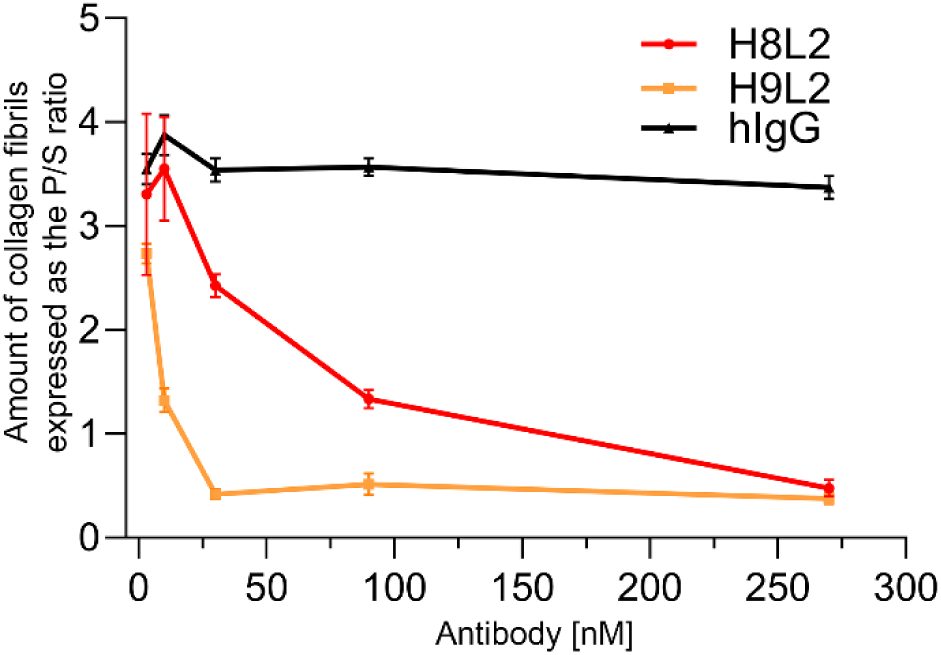
Dose-dependent inhibition of collagen fibril formation by antibodies H8L2 and H9L2. Fibrillogenesis was evaluated across varying concentrations of inhibitory (H8L2, H9L2) and non-inhibitory (hIgG) antibodies. Following fibril formation, samples were fractionated by centrifugation into a fibrillar pellet (P) and a non-aggregated monomeric supernatant (S) for electrophoretic analysis (Fig. 7). Fibril yield is quantified as the ratio of fibrillar to monomeric collagen (P/S ratio). Data represent the mean ± SD of three technical replicates.

These electrophoretic results were further corroborated by microscopic assays, which visually confirmed that the H8L2 and H9L2 variants, but not the hIgG control, effectively prevented the formation of collagen fibrils (Fig. 9).

**Figure 9.**
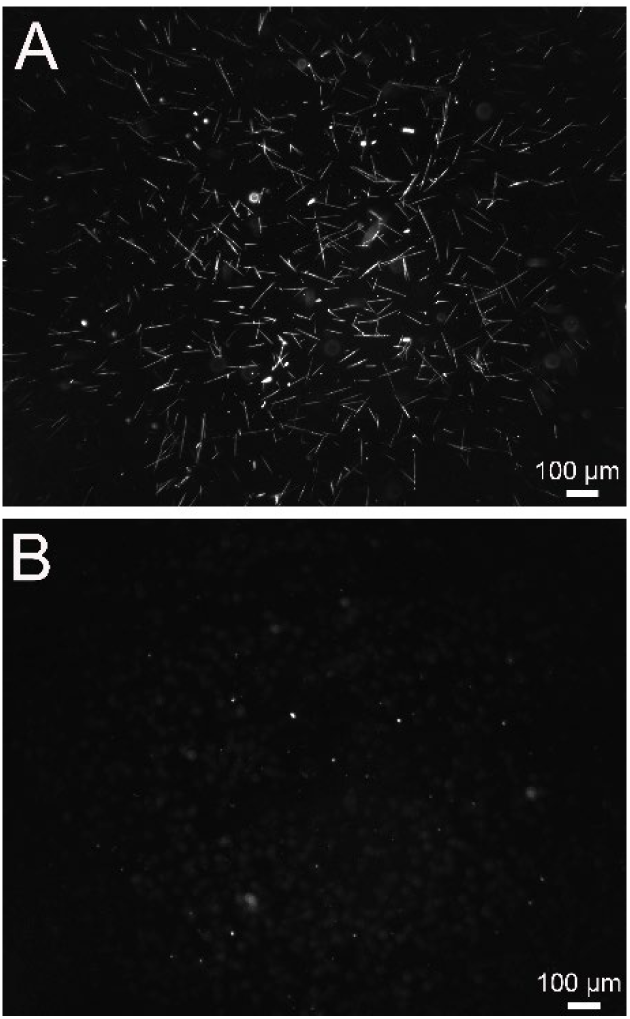
Representative images of collagen fibrils formed in vitro in the presence of inactive human IgG (A) or the presence of the H8L2 or H9L2 variant (B). Bar = 100 μm. In the absence of inhibitory antibodies, numerous fibrils form; in their presence, fibril formation is blocked.

### Analyses of third-generation cbpACA constructs select a candidate with optimal target binding and tissue retention properties

As the final step in optimizing the ACA, we engineered three variants that included the CBP domain at three rationally selected locations, as indicated in Fig. 2.

As with other variants, the cbpACA constructs exhibited the expected electrophoretic migration of the heavy and light chains, as well as chromatographic elution profiles (data not shown).

#### Biosensor-based assays of the cbpACA variants’ binding kinetics

Since the primary reason for engineering the cbpACA variants was to enhance their retention at collagen-rich target sites, we analyzed their binding kinetics. Then, we compared them with those of an unmodified, parent H9L2 ACA variant. The selected binding targets included intact rabbit procollagen I, rabbit atelo-collagen I generated by pepsin digestion of procollagen I, collagen extracted from rabbit tendon using pepsin, mouse skin-derived collagen extracted using acetic acid, collagen II from chick sternum, and synthetic α2Ct peptide. Table IV presents the kinetics of binding of the C-cbpACA, U-cbpACA, and D-cbpACA variants, and the unmodified H9L2 antibody construct, to the above targets. As expected, all antibody variants showed high affinity for collagen variants bearing the α2Ct target. The antibody variants with CBP domains, however, showed markedly higher affinities for collagen targets lacking the canonical α2Ct epitope (i.e., mouse collagen I, chick collagen II, atelo-collagen I, and tendon collagen). All three CBP-containing variants showed similar collagen-binding affinities, though they differed in the k*_a_* and k*_d_* parameters.

**Table. IV.** Binding kinetics of ACA variants containing the CBP cassette.

| Target | ACA variant | $k_{on} [\text{M}^{-1}\text{s}^{-1}]$ | $k_{off} [\text{s}^{-1}]$ | $K_D [\text{M}]$ |
| --- | --- | --- | --- | --- |
| Rabbit Pro-I | ch1ACA (no CBP) | 1.20E+05 | 2.40E-04 | 2.10E-09 |
|  | H9L2 (no CBP) | 9.10E+05 | 1.50E-02 | 1.60E-08 |
|  | U-cbpACA | 6.50E+05 | 3.70E-02 | 5.60E-08 |
|  | D-cbpACA | 2.00E+05 | 1.10E-02 | 5.20E-08 |
|  | C-cbpACA | 4.30E+07 | 5.60E-02 | 1.30E-09 |
| Rabbit Pro-I-peps | ch1ACA (no CBP) | 1.90E+04 | 9.30E-03 | 4.80E-07 |
|  | H9L2 (no CBP) | 1.50E+06 | 2.30E-02 | 1.50E-08 |
|  | U-cbpACA | 3.30E+05 | 2.40E-02 | 7.10E-08 |
|  | D-cbpACA | 9.60E+05 | 9.40E-03 | 9.80E-09 |
|  | C-cbpACA | 2.60E+05 | 3.10E-03 | 1.20E-08 |
| Rabbit tendon collagen I | ch1ACA (no CBP) | 1.20E+04 | 1.93E-02 | 1.60E-06 |
|  | H9L2 (no CBP) | 4.50E+05 | 1.30E-01 | 2.90E-07 |
|  | U-cbpACA | 5.40E+05 | 3.20E-02 | 5.90E-08 |
|  | D-cbpACA | 3.10E+05 | 1.50E-02 | 5.00E-08 |
|  | C-cbpACA | 2.60E+04 | 2.30E-02 | 9.00E-07 |
| Mouse skin collagen I | ch1ACA (no CBP) | 2.60E+04 | 2.50E-03 | 1.30E-06 |
|  | H9L2 (no CBP) | 5.20E+03 | 1.01 | 1.90E-04 |
|  | U-cbpACA | 8.50E+05 | 3.20E-02 | 3.70E-08 |
|  | D-cbpACA | 7.10E+05 | 9.90E-03 | 1.40E-08 |
|  | C-cbpACA | 2.60E+06 | 2.70E-03 | 1.10E-09 |
| Chick sternum collagen II | ch1ACA (no CBP) | 7.00E+03 | 2.10E-02 | 2.90E-06 |
|  | H9L2 (no CBP) | 4.40E+05 | 8.30E-01 | 1.90E-06 |
|  | U-cbpACA | 1.10E+06 | 3.40E-02 | 3.00E-08 |
|  | D-cbpACA | 3.00E+05 | 1.60E-02 | 5.20E-08 |
|  | C-cbpACA | 2.40E+05 | 1.80E-02 | 7.30E-08 |
| $\alpha 2\text{Ct}$ peptide | ch1ACA (no CBP) | 5.90E+04 | 3.60E-06 | 6.10E-11 |
|  | H9L2 (no CBP) | 5.40E+04 | <1E-7 | <1E-12 |
|  | U-cbpACA | 5.60E+04 | 8.10E-05 | 1.50E-09 |
|  | D-cbpACA | 5.50E+04 | <1E-7 | <1E-12 |
|  | C-cbpACA | 6.80E+04 | <1E-7 | <1E-12 |

#### Retention of the C-cbpACA variant in cell culture conditions

We analyzed retention of the C-cbpACA variant within the collagen-rich matrix formed by cultured rabbit dermal fibroblasts (Fig. 10). Using Western blot, we showed that, in contrast to the ACA variant lacking the CBP cassette, the presence of the CBP cassette markedly increased the amount of C-cbpACA in the matrix (Fig. 11).

**Figure 10.**
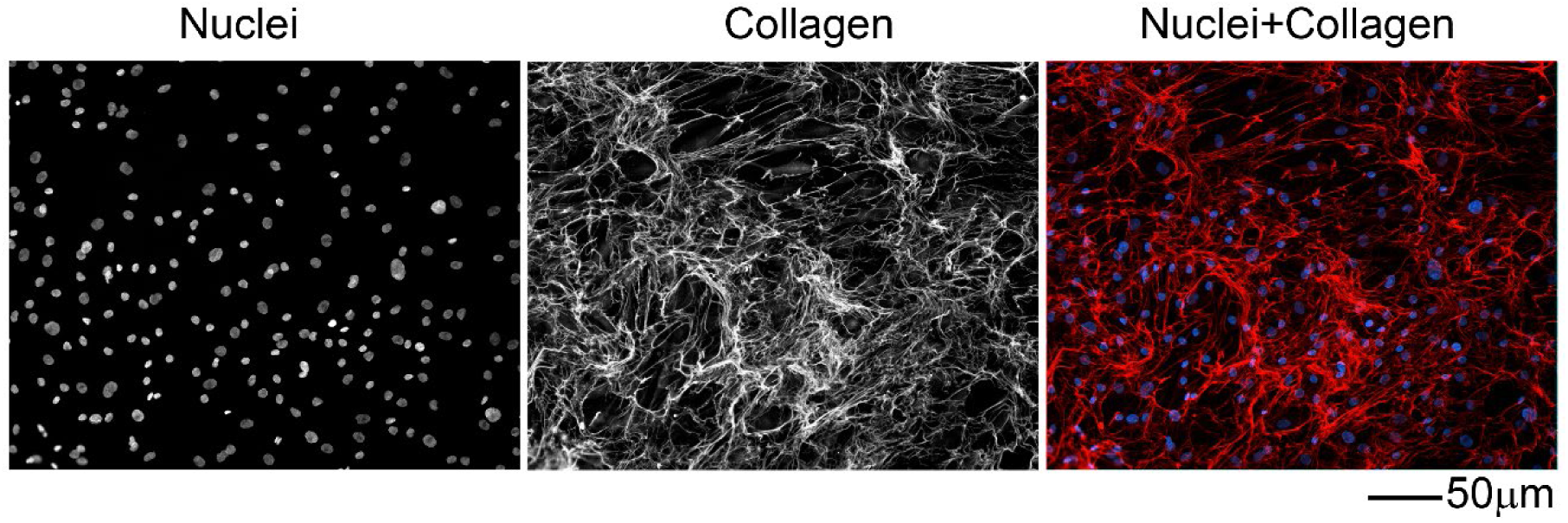
Visualization of the collagen-rich extracellular matrix produced by cultured fibroblasts. Red signals represent collagen fibrils detected with specific antibodies. Nuclei are labeled blue with DAPI to provide a cellular context.

**Figure 11.**
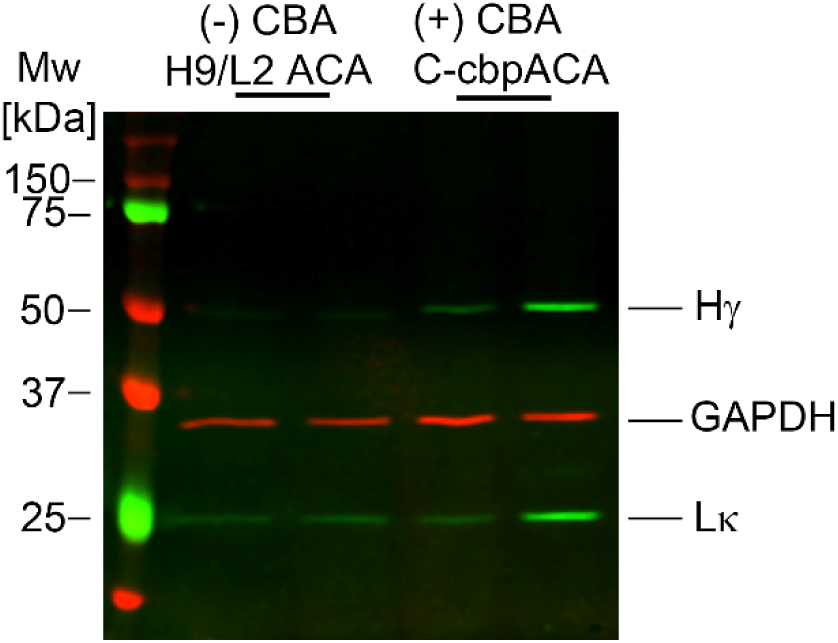
Western blot analysis of matrix retention for unmodified ACA and the C-cbpACA variants. Proteins were extracted from the collagen-rich cell layers described in Figure 10. The heavy (Hγ) and light (Lκ) chains of ACA, alongside glyceraldehyde 3-phosphate dehydrogenase (GAPDH), are indicated.

#### Assay of acute and chronic cytotoxicity

Employing rabbit dermal fibroblasts and human keloid-derived fibroblasts, our studies demonstrated that C-cbpACA did not negatively impact the viability or proliferation of these cells. In contrast, actinomycin D, used as a positive control, markedly inhibited fibroblast proliferation (Figs. S8 and S9).

### Assays of the C-cbpACA construct demonstrate its anti-fibrotic efficacy and safety in the rabbit arthrofibrosis model

#### Mechanical assays of the flexion contracture

A Kruskal-Wallis H test was conducted to determine if there were differences in the flexion contracture scores (defined as uninjured knee angle/injured knee angle ratio measured at 0.2 Nm torque) among three groups: an untreated CTR group, a group treated with standard ACA (ACA), and a pilot group treated with C-cbpACA.

Visual inspection of the boxplots indicated higher median scores and greater variance in the untreated CTR group than in either treatment group. The test revealed a significant difference in flexion contracture scores across the three categories (H(2) = 9.319, p = 0.009), with a total sample size of 28. Pairwise comparisons with a Bonferroni correction revealed significant reductions in flexion contracture for both the standard ACA treatment (p = 0.027) and the C-cbpACA pilot treatment (p = 0.043) when compared to the untreated CTR group. However, the analysis showed no significant difference in flexion contracture between the standard ACA and C-cbpACA treatment groups (p = 1.000; Fig. 12).

**Figure 12.**
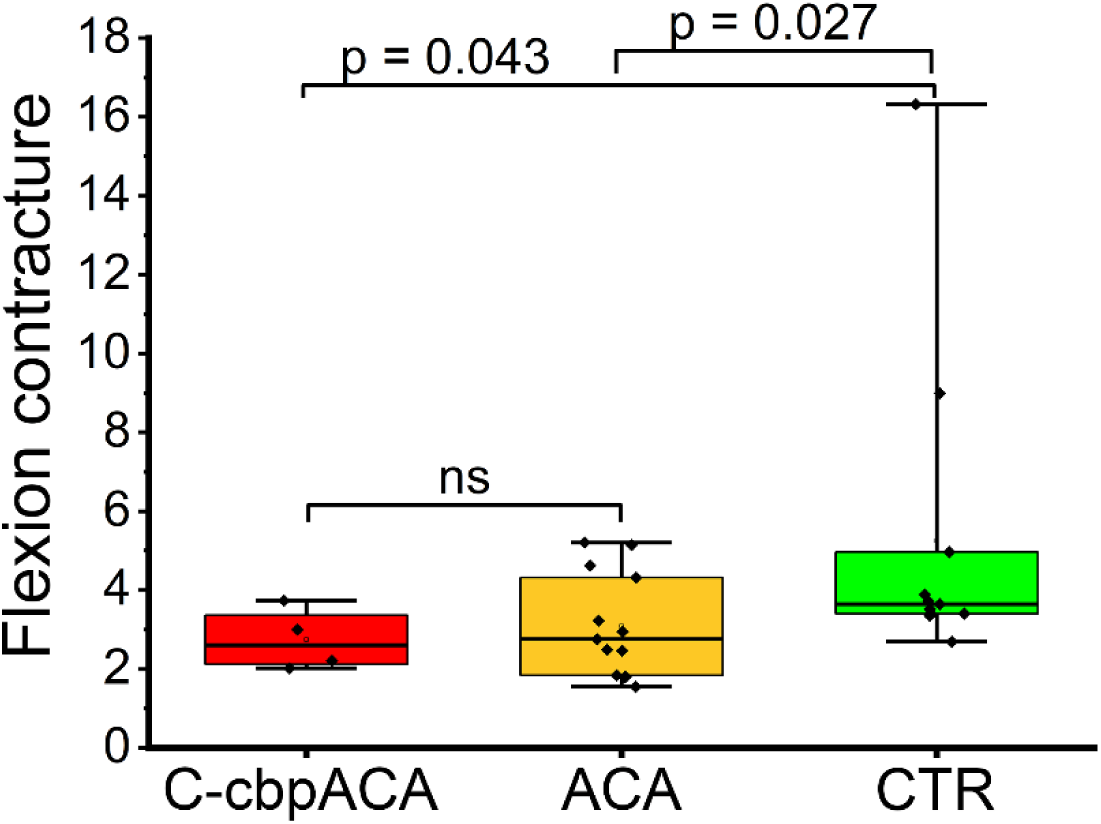
Flexion contracture following knee injury. Data are expressed as the ratio of maximal knee extension angles in uninjured versus injured knees at 0.2 Nm torque. Boxes represent the interquartile range (25th to 75th percentiles), horizontal lines indicate medians, and whiskers denote minimum and maximum values.

#### Relative collagen content

A Kruskal-Wallis H test evaluated differences in relative collagen content among an untreated CTR, a standard ACA treatment group, and a pilot group treated with C-cbpACA (Fig. 13). The analysis used a total sample size of 24, with a notably unbalanced design due to the pilot nature of the C-cbpACA group (n = 4).

**Figure 13.**
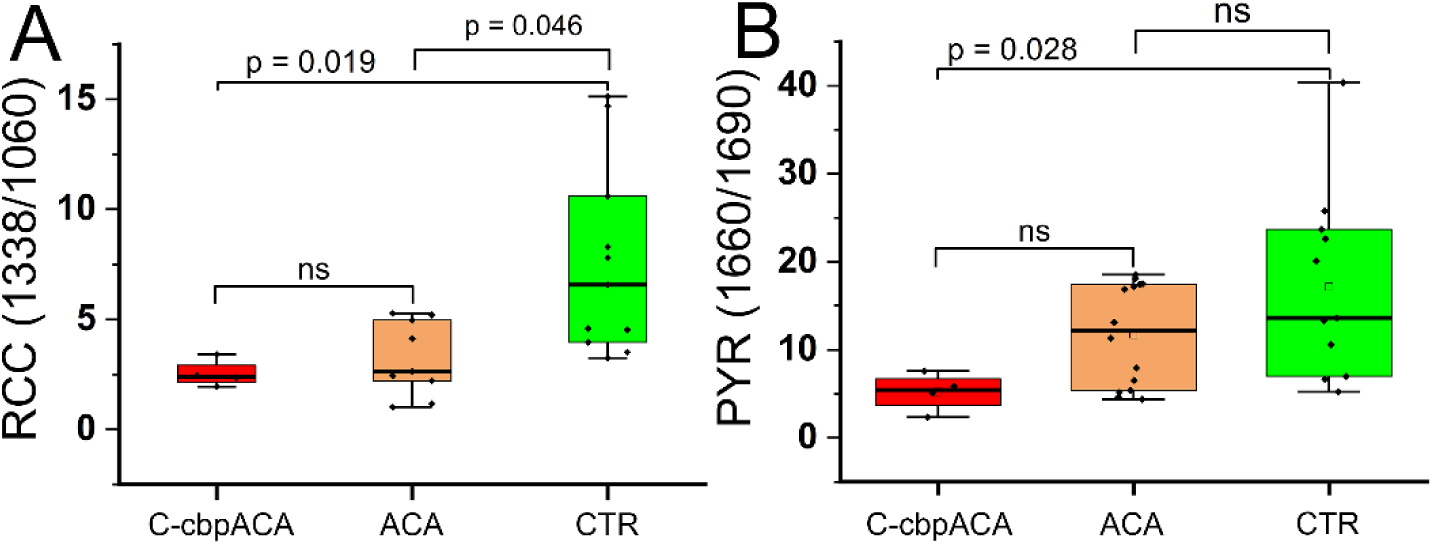
FTIR analysis of collagen content and cross-link maturity in posterior capsule scar tissue. (A) Relative amount of collagen (RCC) and (B) maturity of collagen cross-links (PYR) in tissue from C-cbpACA-treated, unmodified ACA-treated, and control (CTR) rabbits. RCC is calculated as the ratio of spectral peak areas for collagen (1338 cm⁻¹ ν) to sulfated GAGs (1064 cm⁻¹ ν). PYR cross-link maturity is defined as the ratio of peak areas for mature trivalent PYR (1660 cm⁻¹ ν) to immature divalent deDHLNL cross-links (1690 cm⁻¹ ν). Boxes represent the interquartile range with the median indicated by a horizontal line; whiskers denote minimum and maximum values.

The test revealed a significant difference in collagen content across the three groups (H(2) = 9.952, p = 0.007). Post-hoc pairwise comparisons with a Bonferroni correction for multiple testing revealed significant reductions in collagen content for both the standard ACA treatment (p = 0.046) and the modified C-cbpACA pilot treatment (p = 0.019) when compared to the untreated CTR group. The analysis showed no significant difference between the standard ACA and C-cbpACA treatments (p = 1.000).

#### Relative PYR content

A Kruskal-Wallis H test was used to evaluate differences in relative PYR content among the three groups (Fig. 13). The analysis included a total of 29 samples, with a notably unbalanced design due to the C-cbpACA group (n = 4). The test revealed a significant difference in relative PYR content across the three groups (H(2) = 6.997, p = 0.030). Post-hoc pairwise comparisons with a Bonferroni correction revealed a significant reduction in relative PYR content in the C-cbpACA pilot treatment group (p = 0.028) compared with the untreated CTR group. Conversely, the standard ACA treatment did not show a significant difference compared with the untreated CTR group (p = 0.443). When comparing the two active antibodies, the analysis showed no significant difference between the standard ACA and modified ACA treatments (p = 0.297).

#### Total collagen content (Col-Un/In)

A Kruskal-Wallis H test was used to evaluate differences in total collagen content (as measured by hydroxyproline assays) among the three groups.

First, total collagen content was expressed as the amount of this protein per unit of dry PC tissue mass. Subsequently, the uninjured knee-to-injured knee ratios (Col-Un/In) were calculated. Hence, the higher the Col-Un/In ratio, the lower the collagen content in PCs from injured knees.

The analysis utilized a total sample size of 43, including the unbalanced preliminary C-cbpACA pilot group (n = 4). The test revealed a significant difference in Col-Un/In ratios across the three categories (H(2) = 6.294, p = 0.043). Post-hoc pairwise comparisons with a Bonferroni correction revealed significant differences in the ratios for the C-cbpACA pilot treatment compared to both the standard ACA treatment (p = 0.048) and the untreated CTR group (p = 0.050).

Visual inspection of the data distributions indicated that the modified antibody formulation yielded a higher median ratio than the other groups. Conversely, there was no statistically significant difference between the standard ACA treatment group and the untreated CTR group (p = 1.000; Fig. 14).

**Figure 14.**
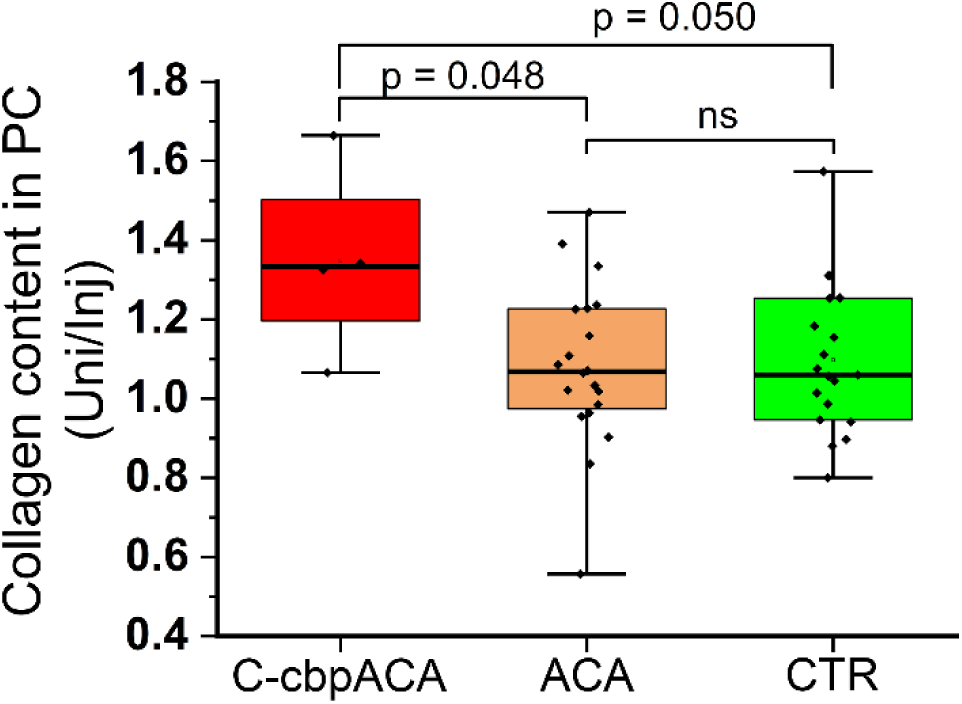
Collagen content in posterior capsules. Collagen was quantified by measuring hydroxyproline levels. Results are expressed as the ratio of collagen in uninjured versus injured capsules. Boxes represent the interquartile range with the median indicated by a horizontal line; whiskers denote minimum and maximum values.

While these findings must still be interpreted with caution due to the pilot group’s exceptionally small sample size, this is a particularly notable outcome: it represents a distinct statistical divergence between the two active formulations, with the modified antibody driving a measurable shift in hydroxyproline levels that the standard formulation did not.

#### Histological assays of collagen fibrils

In contrast to uninjured knees, H&E staining of the PCs isolated from injured knees demonstrated infiltration of inflammatory and fibroblastic cells and deposition of fibrotic tissue (Fig. 15). Fibrotic tissue formed in the injured PCs was characterized by a marked increase in thin, GB fibrils Fig. 15 and 16). Quantification of the percentage of RB, YB, and GB fibrils suggested that, in comparison to CTR, unmodified ACA and the C-cbpACA variant markedly decreased the percentage of the GB fibrils fraction observed in the injured PCs (Fig. 16). In uninjured PCs, the percentages of thick fibrils represented by RB and YB fractions as well as thin fibrils represented by the GB fraction were comparable (Fig. 16).

**Figure 15.**
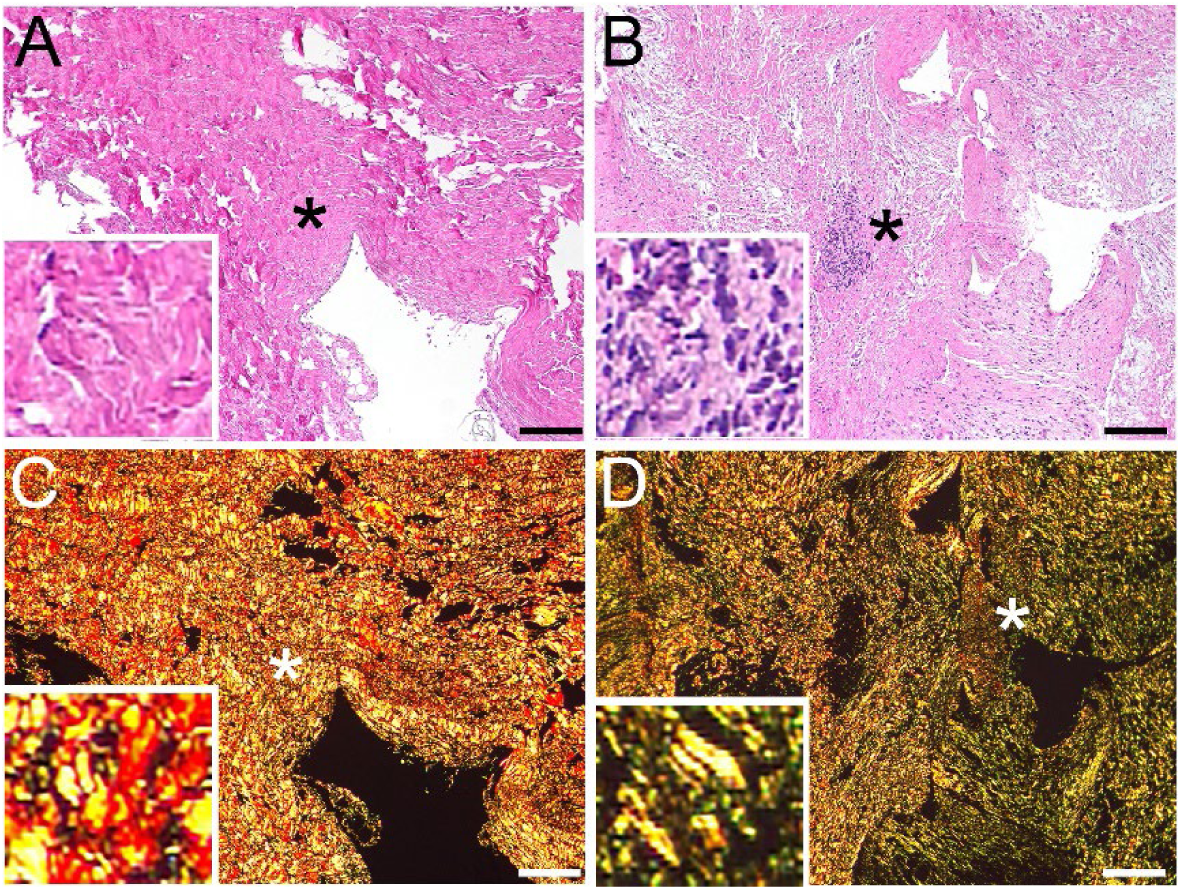
Microscopic analysis of posterior capsules (PCs) from uninjured and injured knees. (A, B) H&E staining reveals sparse cellularity in the uninjured PC (A), compared to a marked invasion of fibroblastic cells in the injured PC (B). (C, D) Picrosirius red staining demonstrates a shift in collagen birefringence, from predominantly red/yellow in the uninjured PC (C) to green in the injured PC (D). Insets highlight high-magnification details of the areas marked with asterisks (*). Bars = 100 µm

**Figure 16.**
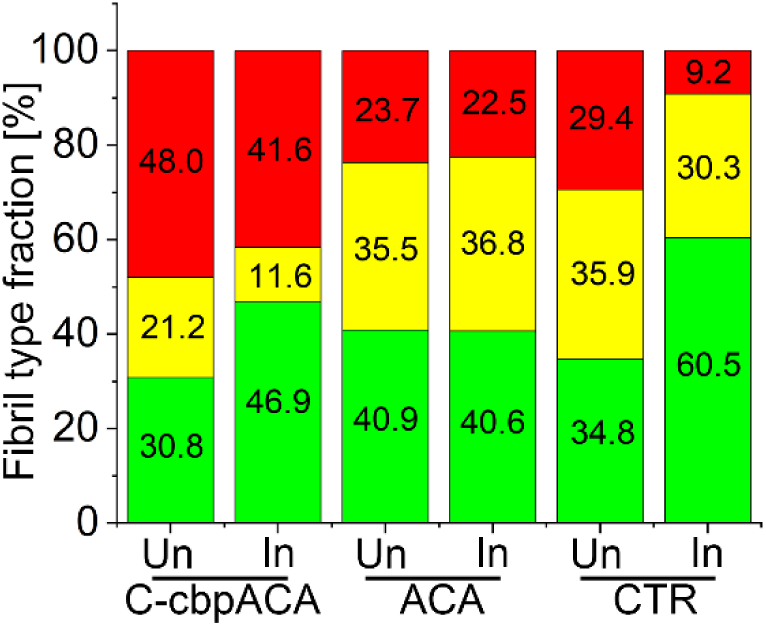
Quantification of collagen fibril populations in uninjured (Un) and injured (In) posterior capsules (PCs). Fibril subtypes in tissues from C-cbpACA-treated, unmodified ACA-treated, and control (CTR) rabbits were evaluated using picrosirius red staining. Populations are expressed as the percentage of fibrils exhibiting red, yellow, or green birefringence under polarized light.

#### Complete blood count

CBC provides valuable information about potential pathological changes that could have occurred due to the C-cbpACA application. As indicated in Fig. S10 and Tabs. Va and Vb, all cellular parameters stayed within their respective rabbit-specific physiological ranges.

**Table Va.** CBC in the C-cbpACA-treated rabbits.

| Time<br>[weeks] | Rabbit | Blood parameter |  |  |  |  |  |  |  |  |  |
| --- | --- | --- | --- | --- | --- | --- | --- | --- | --- | --- | --- |
|  |  | WBC | NEU | LYM | MONO | EOS | BAS | NEU % | LYM % | MONO % | EOS % |
| 0 | 1 | 4.59 | 0.95 | 3.31 | 0.26 | 0.03 | 0.04 | 20.6 | 72 | 5.7 | 0.8 |
|  | 2 | 2.13 | 0.61 | 1.44 | 0.06 | 0.01 | 0.01 | 28.3 | 67.4 | 3 | 0.6 |
|  | 3 | 5.9 | 0.79 | 4.61 | 0.4 | 0.04 | 0.06 | 13.3 | 78.1 | 6.9 | 0.7 |
|  | 4 | 5.14 | 1.36 | 3.32 | 0.34 | 0.05 | 0.07 | 26.5 | 64.6 | 6.4 | 1.1 |
| 4 | 1 | 5.54 | 0.82 | 4.19 | 0.38 | 0.1 | 0.05 | 14.8 | 75.6 | 6.8 | 1.9 |
|  | 2 | 7.15 | 1.11 | 5.23 | 0.53 | 0.16 | 0.12 | 15.5 | 73.1 | 7.4 | 2.3 |
|  | 3 | 7.66 | 2.22 | 4.23 | 0.92 | 0.07 | 0.22 | 29 | 55.2 | 12.1 | 0.9 |
|  | 4 | 6.84 | 1.08 | 4.67 | 0.66 | 0.38 | 0.05 | 15.8 | 68.3 | 9.6 | 5.5 |
| 8 | 1 | 4.7 | 1.26 | 2.58 | 0.62 | 0.09 | 0.15 | 26.8 | 54.8 | 13.1 | 2 |
|  | 2 | 5.51 | 0.73 | 4.39 | 0.31 | 0.07 | 0.01 | 13.2 | 79.7 | 5.5 | 1.3 |
|  | 3 | 5.91 | 1.6 | 3.55 | 0.63 | 0.05 | 0.08 | 26.9 | 60 | 10.8 | 0.9 |
|  | 4 | 6.76 | 0.8 | 5.44 | 0.44 | 0.02 | 0.06 | 11.8 | 80.4 | 6.6 | 0.3 |
| 12 | 1 | 5.07 | 0.56 | 4.04 | 0.38 | 0.07 | 0.02 | 10.9 | 79.7 | 7.5 | 1.5 |
|  | 2 | 3.8 | 0.44 | 2.97 | 0.28 | 0.03 | 0.08 | 11.6 | 78.1 | 7.3 | 0.8 |
|  | 3 | 6.66 | 0.67 | 5.56 | 0.28 | 0.05 | 0.1 | 10.1 | 83.5 | 4.1 | 0.8 |
|  | 4 | 3.07 | 0.52 | 2.33 | 0.18 | 0.02 | 0.02 | 16.8 | 75.9 | 5.8 | 0.7 |
| Normal range |  | 3.0-13.50<br>[10<sup>3</sup>/μl] | 0.5-6.6<br>[10<sup>3</sup>/μl] | 1.0-6.8<br>[10<sup>3</sup>/μl] | 0.08-1.51<br>[10<sup>3</sup>/μl] | 0.0-0.51<br>[10<sup>3</sup>/μl] | 0.0-0.76<br>[10<sup>3</sup>/μl] | 14.0-62.0<br>[%] | 25.0-82.0<br>[%] | 2.0-15.0<br>[%] | 0.0-6.0<br>[%] |

WBC; white blood cells, NEU; neutrophils, LYM; lymphocytes, MONO; monocytes, EOS; eosinophils, BAS; basophils, NEU %; percentage of neutrophils, LYM %; percentage of lymphocytes, MONO %; percentage of monocytes, EOS %; percentage of eosinophils

**Table Vb:** CBC in the C-cbpACA-treated rabbits.

| Time [weeks] | Rabbit | Blood parameter |  |  |  |  |  |  |  |  |  |
| --- | --- | --- | --- | --- | --- | --- | --- | --- | --- | --- | --- |
|  |  | BAS % | HCT | HGB | RBC | MCV | RDW % | MCH | MCHC | PLT | MPV |
| 0 | 1 | 0.9 | 30.6 | 10.7 | 5.5 | 55.7 | 14.3 | 19.5 | 35.1 | 185 | 5.9 |
|  | 2 | 0.7 | 29.5 | 10.4 | 4.71 | 62.5 | 12.8 | 22 | 35.2 | 247 | 6.8 |
|  | 3 | 1 | 35.7 | 12.4 | 5.66 | 63 | 11.9 | 21.8 | 34.7 | 375 | 5 |
|  | 4 | 1.4 | 34.8 | 11.8 | 5.24 | 66.4 | 13.3 | 22.6 | 34 | 380 | 5.6 |
| 4 | 1 | 0.9 | 33.6 | 12.0 | 5.52 | 60.8 | 12 | 21.7 | 35.6 | 463 | 5.8 |
|  | 2 | 1.7 | 36.4 | 12.9 | 6.41 | 56.9 | 11.8 | 20.1 | 35.4 | 432 | 5.3 |
|  | 3 | 2.8 | 31 | 11.2 | 5.02 | 61.8 | 11.8 | 22.3 | 36.1 | 851 | 4.9 |
|  | 4 | 0.8 | 32.9 | 11.5 | 5.3 | 62 | 13.2 | 21.8 | 35.1 | 774 | 4.8 |
| 8 | 1 | 3.3 | 31.6 | 11 | 5.6 | 56.4 | 12.2 | 19.7 | 35 | 379 | 5.5 |
|  | 2 | 0.3 | 33.1 | 11.6 | 5.59 | 59.2 | 13.3 | 20.8 | 35.2 | 466 | 5.9 |
|  | 3 | 1.4 | 29.8 | 10.5 | 4.76 | 62.6 | 12.3 | 22 | 35.2 | 949 | 4.9 |
|  | 4 | 0.9 | 32.4 | 11.5 | 5.48 | 59.1 | 13.6 | 20.9 | 35.4 | 971 | 4.4 |
| 12 | 1 | 0.4 | 32.2 | 11 | 5.44 | 59.2 | 14.1 | 20.3 | 34.3 | 519 | 6.1 |
|  | 2 | 2.2 | 33.7 | 11.5 | 5.84 | 57.7 | 13.7 | 19.7 | 34.1 | 333 | 5.4 |
|  | 3 | 1.5 | 35.4 | 12 | 5.91 | 60 | 13.9 | 20.4 | 34 | 647 | 5 |
|  | 4 | 0.8 | 29.1 | 10.1 | 4.8 | 60.7 | 13.8 | 21.1 | 34.7 | 166 | 5.4 |
| Normal range |  | 0.0-8.0 [%] | 25.0-42.0 [%] | 8.0-14 [g/dL] | 3.40-6.50 [10<sup>6</sup>/μl] | 60.0-80.0 [fL] | 10.5-22.0 [%] | 19.0-25.0 [pg] | 30.0-36.0 [g/dL] | 100-1250 [10<sup>3</sup>/μl] | 4.0-7.8 [fL] |

BAS %; percentage of basophils, HCT; hematocrit, HGB; hemoglobin, RBC; red blood cells, MCV; mean corpuscular volume, RDW %; red blood cell distribution width, MCH; mean corpuscular hemoglobin, MCHC; mean corpuscular hemoglobin concentration, PLT; platelet count, MPV; mean platelet volume.

#### Histology of tissues and organs

Because of the dynamic exchange of macromolecules between the intraarticular space and the lymphatic and circulatory systems, we assessed whether continuous administration of the C-cbpACA had any adverse effects on distant tissues and organs. Histology of crucial tissues and organs collected from the rabbits did not show any structural abnormalities or excess inflammatory cells. Supplementary figures present histological images of the assayed specimens (Figs. S11-S15).

## DISCUSSION

The development of safe and effective anti-fibrotic therapies remains a major unmet clinical need. The strategy described here targets the assembly of collagen I molecules into insoluble fibrils. By interfering with the precise molecular interaction between α2Ct and the TBR, we selectively attenuated pathological collagen accumulation while preserving upstream cellular processes essential for normal tissue repair (2).

A central achievement of this work is the successful humanization and optimization of a prototypic ACA without compromising target specificity or functional activity. Humanization of murine antibodies often results in reduced affinity or altered binding kinetics. Here, however, a structure-guided approach using three-dimensional homology modeling enabled rational selection of a framework and targeted back-mutation of key murine residues. This strategy preserved the spatial architecture of the antigen-binding site while simultaneously improving the antibody’s chemical stability and developability by eliminating deamidation-prone motifs. The resulting lead variants, H8L2 and H9L2, demonstrated excellent structural integrity, homogeneity, and robust expression in a clinically relevant mammalian production system.

Importantly, biochemical and biophysical analyses confirmed that these optimizations did not compromise the antibody’s mechanism of action. Biosensor-based studies demonstrated high-affinity binding of the humanized variants not only to a synthetic α2Ct peptide but also to native, intact procollagen I, underscoring the physiological relevance of the interaction. Functional assays further showed that these antibodies effectively blocked de novo collagen fibrillogenesis in vitro, trapping collagen molecules in a non-aggregated monomeric state. These effects were consistently confirmed using complementary electrophoretic and microscopic approaches, strengthening confidence in the robustness of the observed inhibitory activity.

To address the translational challenge of achieving effective and durable localization within collagen-rich fibrotic tissues, we engineered a third-generation ACA construct incorporating a CBP. The rationale for this modification was informed by prior successful applications of the TKKTLRT peptide to enhance matrix retention of diverse biologics (47–50). By conferring dual binding functionality (i.e., targeting both the α2Ct epitope and the triple-helical region of collagen I), the CBP-modified constructs were designed to increase residence time at sites of active fibrosis without altering intrinsic antigen specificity.

Among the CBP-containing variants, fusion of the peptide to the C-cbpACA emerged as the optimal configuration. Biosensor analyses revealed that this construct retained, and in fact enhanced, binding affinity for procollagen I relative to its unmodified parent antibody. The nanomolar dissociation constant observed for C-cbpACA reflects a favorable combination of rapid association and slow dissociation kinetics, consistent with stable retention in collagen-rich matrices. Cell-based assays corroborated these findings, demonstrating markedly increased retention of the CBP-modified antibody within fibroblast-derived collagen matrices under conditions of repeated media exchange.

The biological relevance of this enhanced targeting was supported by results obtained in a rabbit model of post-traumatic arthrofibrosis. Although conducted as a pilot study, local administration of C-cbpACA resulted in measurable improvements across multiple independent outcome metrics. These included reductions in joint flexion contracture, decreases in total and relative collagen content, lower levels of mature pyridinoline cross-links, and a shift in collagen fibril architecture away from thin, densely deposited fibrils associated with active fibrosis. Notably, these changes were observed without detectable systemic toxicity, as evidenced by normal CBC and unremarkable histological findings in distant organs. These results are consistent with the CBC and histological assays we performed earlier in rabbits treated with unmodified ACA (2).

While direct comparisons between the modified and unmodified ACA formulations were limited by the small size of the pilot cohort, several endpoints, particularly hydroxyproline-based collagen quantification, suggest a potentially enhanced anti-fibrotic effect of the CBP-modified construct. These findings are best interpreted as supportive evidence that targeted matrix retention can augment the functional impact of extracellular collagen assembly inhibitors.

In sum, this study established a comprehensive framework for the rational engineering of clinically viable anti-fibrotic antibodies that act extracellularly and exhibit enhanced localization within fibrotic tissues. By combining precise molecular targeting of collagen fibrillogenesis with engineered matrix affinity, the C-cbpACA construct represents a promising therapeutic strategy for conditions characterized by pathological scarring. The results presented here justify further, fully powered preclinical studies to optimize dosing, delivery, and long-term efficacy, and to explore the applicability of this approach across a broader spectrum of fibrotic diseases.

## LIMITATIONS

Several limitations of the present study should be acknowledged, particularly given the pilot nature of the in vivo investigations. While the engineering, biochemical, and cell-based evaluations of the humanized ACA variants were extensive and comprehensive, the animal studies were intentionally designed as an exploratory, hypothesis-generating assessment to determine whether the CBP–modified ACA warranted further, fully powered preclinical investigation.

First, the rabbit arthrofibrosis experiments involving C-cbpACA had an exceptionally small sample size. Only four animals were included in the C-cbpACA treatment group, reflecting both ethical considerations to minimize animal use and the study’s preliminary nature. As a result, the statistical power to detect differences, particularly between the modified C-cbpACA and the standard ACA formulation, was inherently limited. Although statistically significant differences relative to untreated controls were observed in several biomechanical, biochemical, and spectroscopic endpoints, comparisons between the two active antibody formulations must be interpreted with caution. The absence of statistically significant differences in some direct comparisons is therefore more likely attributable to insufficient power rather than equivalence of efficacy.

Second, the animal study relied in part on historical control cohorts, including untreated controls and rabbits previously treated with unmodified ACA variants. Although these cohorts were generated using identical surgical procedures, outcome measures, and analytical pipelines, the use of historical controls introduces unavoidable sources of variability that cannot be fully controlled in a single pilot experiment. This design choice, however, was deliberate and ethically motivated, allowing meaningful contextual interpretation of the C-cbpACA outcomes while reducing redundancy and limiting animal use. Nonetheless, future studies incorporating concurrent randomized control groups will be essential to rigorously quantify the incremental benefit of the CBP modification.

Third, the study focused on a single dosing strategy, delivery route, and treatment duration. While localized intra-articular delivery via an implanted pump is well-suited for mechanistic proof-of-concept studies, it does not capture the full translational landscape of potential clinical applications. Dose response relationships, alternative delivery regimens, temporal windows of intervention, and long-term durability of the anti-fibrotic effects were beyond the scope of this pilot investigation and remain important areas for future work.

Finally, although the rabbit model of arthrofibrosis is highly relevant and well validated, particularly given the complete conservation of the α2Ct epitope between rabbits and humans, it represents one specific fibrotic context. Excessive scarring occurs across diverse tissues and pathological conditions, each with distinct biomechanical environments and inflammatory drivers. Consequently, the generalizability of the observed effects of C-cbpACA to other fibrotic indications must be evaluated in additional disease models.

Taken together, these limitations reflect intentional design choices appropriate for a pilot study, whose primary objective was to establish the feasibility, biological activity, and safety of a rationally engineered, matrix-targeted ACA. The collective findings strongly justify advancing to larger, prospectively powered animal studies to more definitively assess efficacy, optimize dosing strategies, and refine translational potential.

### NOTE

Although we provided essential information regarding the engineering of the humanized constructs above, their detailed sequences are not included due to a pending patent application. The PCT application was filed on January 12, 2026 (Attorney Docket No.: 205961-7120WO1(00553)).

## ACKNOWLEDGEMENTS

The authors are grateful to veterinarians and animal health staff for excellent veterinary assistance. The authors wish to acknowledge the late Dr. Andrzej Steplewski for his contribution.

## AUTHOR CONTRIBUTIONS

Conceptualization: AM, BY, JL, AF Funding acquisition: AM, JL, AF Investigation: AM, BY, JL, AF, JF Supervision: AM, BY, JL, AF Formal analysis: AM, BY, JL, AF, JF Writing original draft: AF Review and editing: AM, BY, JL, JF Administration: JL, AF Visualization: AM, BY, JL, AF, JF Methodology: AM, BY, JL, AF, JF

## REFERENCES

1. Wynn TA. Cellular and molecular mechanisms of fibrosis. The Journal of pathology. 2008;214(2):199–210.

2. Steplewski A, Fertala J, Tomlinson RE, Wang ML, Donahue A, Arnold WV, et al. Mechanisms of reducing joint stiffness by blocking collagen fibrillogenesis in a rabbit model of posttraumatic arthrofibrosis. PLoS One. 2021;16(9):e0257147.

3. Chen AF, Lee YS, Seidl AJ, Abboud JA. Arthrofibrosis and large joint scarring. Connect Tissue Res. 2019;60(1):21–8.

4. Fertala J, Rivlin M, Wang ML, Beredjiklian PK, Steplewski A, Fertala A. Collagen-rich deposit formation in the sciatic nerve after injury and surgical repair: A study of collagen-producing cells in a rabbit model. Brain Behav. 2020;10(10):e01802.

5. Wang ML, Rivlin M, Graham JG, Beredjiklian PK. Peripheral nerve injury, scarring, and recovery. Connect Tissue Res. 2018:1–7.

6. Steplewski A, Fertala J, Beredjiklian PK, Abboud JA, Wang ML, Namdari S, et al. Auxiliary proteins that facilitate formation of collagen-rich deposits in the posterior knee capsule in a rabbit-based joint contracture model. J Orthop Res. 2016;34(3):489–501.

7. Friedman SL, Sheppard D, Duffield JS, Violette S. Therapy for fibrotic diseases: nearing the starting line. Science translational medicine. 2013;5(167):167sr1.

8. Chung HJ, Steplewski A, Chung KY, Uitto J, Fertala A. Collagen fibril formation. A new target to limit fibrosis. J Biol Chem. 2008;283(38):25879–86.

9. Prockop DJ, Fertala A. Inhibition of the self-assembly of collagen I into fibrils with synthetic peptides. Demonstration that assembly is driven by specific binding sites on the monomers. J Biol Chem. 1998;273(25):15598–604.

10. Fertala J, Kostas J, Hou C, Steplewski A, Beredjiklian P, Abboud J, et al. Testing the anti-fibrotic potential of the single-chain Fv antibody against the alpha2 C-terminal telopeptide of collagen I. Connect Tissue Res. 2014;55(2):115–22.

11. Fertala J, Romero F, Summer R, Fertala A. Target-Specific Delivery of an Antibody That Blocks the Formation of Collagen Deposits in Skin and Lung. Monoclonal antibodies in immunodiagnosis and immunotherapy. 2017;36(5):199–207.

12. Fertala J, Steplewski A, Kostas J, Beredjiklian P, Williams G, Arnold W, et al. Engineering and characterization of the chimeric antibody that targets the C-terminal telopeptide of the alpha2 chain of human collagen I: a next step in the quest to reduce localized fibrosis. Connect Tissue Res. 2013;54(3):187–96.

13. Fertala A, Steplewski A, inventors; Thomas Jefferson University, assignee. Engineered antibody for inhibition of fibrosis. USA patent US RE49,477 E. 2023.

14. Steplewski A, Fertala J, Beredjiklian PK, Abboud JA, Wang MLY, Namdari S, et al. Blocking collagen fibril formation in injured knees reduces flexion contracture in a rabbit model. J Orthop Res. 2017;35(5):1038–46.

15. Fertala A, Steplewski A, inventors; Thomas Jefferson University, assignee. Engineered antibody for inhibition of fibrosis USA patent 9,777,055. 2017.

16. Jia L, Sun Y. Protein asparagine deamidation prediction based on structures with machine learning methods. PLoS One. 2017;12(7):e0181347.

17. Yu B, Larrick J, inventors; Larix Bioscience, LLC, assignee. Cell Line Screening Method. patent US 9,910,038 B. 2018.

18. de Souza SJ, Brentani R. Collagen binding site in collagenase can be determined using the concept of sense-antisense peptide interactions. J Biol Chem. 1992;267(19):13763–7.

19. Fiedler-Nagy C, Bruckner P, Hayashi T, Prockop DJ. Isolation of unhydroxylated type I procollagen folding of the protein in vitro. Arch Biochem Biophys. 1981;212(2):668–77.

20. Kadler KE, Hulmes DJ, Hojima Y, Prockop DJ. Assembly of type I collagen fibrils de novo by the specific enzymic cleavage of pC collagen. The fibrils formed at about 37 degrees C are similar in diameter, roundness, and apparent flexibility to the collagen fibrils seen in connective tissue. Ann N Y Acad Sci. 1990;580:214–24.

21. Nesterenko S, Morrey ME, Abdel MP, An KN, Steinmann SP, Morrey BF, et al. New rabbit knee model of posttraumatic joint contracture: indirect capsular damage induces a severe contracture. J Orthop Res. 2009;27(8):1028–32.

22. Barlow JD, Hartzler RU, Abdel MP, Morrey ME, An KN, Steinmann SP, et al. Surgical capsular release reduces flexion contracture in a rabbit model of arthrofibrosis. J Orthop Res. 2013;31(10):1529–32.

23. Hildebrand KA, Sutherland C, Zhang M. Rabbit knee model of post-traumatic joint contractures: the long-term natural history of motion loss and myofibroblasts. J Orthop Res. 2004;22(2):313–20.

24. Hildebrand KA, Holmberg M, Shrive N. A new method to measure post-traumatic joint contractures in the rabbit knee. Journal of biomechanical engineering. 2003;125(6):887–92.

25. Hildebrand KA, Zhang M, Germscheid NM, Wang C, Hart DA. Cellular, matrix, and growth factor components of the joint capsule are modified early in the process of posttraumatic contracture formation in a rabbit model. Acta orthopaedica. 2008;79(1):116–25.

26. Junqueira LC, Bignolas G, Brentani RR. Picrosirius staining plus polarization microscopy, a specific method for collagen detection in tissue sections. The Histochemical journal. 1979;11(4):447–55.

27. Jalil JE, Doering CW, Janicki JS, Pick R, Shroff SG, Weber KT. Fibrillar collagen and myocardial stiffness in the intact hypertrophied rat left ventricle. Circ Res. 1989;64(6):1041–50.

28. Gopinathan PA, Kokila G, Jyothi M, Ananjan C, Pradeep L, Nazir SH. Study of collagen birefringence in different grades of oral squamous cell carcinoma using picrosirius red and polarized light microscopy. Scientifica. 2015;2015:802980-.

29. Junqueira LC, Montes GS, Sanchez EM. The influence of tissue section thickness on the study of collagen by the Picrosirius-polarization method. Histochemistry. 1982;74(1):153–6.

30. Dayan D, Hiss Y, Hirshberg A, Bubis JJ, Wolman M. Are the polarization colors of picrosirius red-stained collagen determined only by the diameter of the fibers? Histochemistry. 1989;93(1):27–9.

31. Montes GS, Nicolelis MA, Brentani-Samaia HP, Furuie SS. Collagen fibril diameters in arteries of mice. A comparison of manual and computer-aided morphometric analyses. Acta Anat (Basel). 1989;135(1):57–61.

32. Allon I, Vered M, Buchner A, Dayan D. Stromal differences in salivary gland tumors of a common histopathogenesis but with different biological behavior: a study with picrosirius red and polarizing microscopy. Acta Histochem. 2006;108(4):259–64.

33. Rich L, Whittaker P. Collagen and picrosirius red staining: A polarized light assessment of fibrillar hue and spatial distribution. Journal of Morphological Sciences. 2005;22(2):97–104.

34. Querido W, Kandel S, Pleshko N. Applications of Vibrational Spectroscopy for Analysis of Connective Tissues. Molecules. 2021;26(4).

35. Cheheltani R, Rosano JM, Wang B, Sabri AK, Pleshko N, Kiani MF. Fourier transform infrared spectroscopic imaging of cardiac tissue to detect collagen deposition after myocardial infarction. J Biomed Opt. 2012;17(5):056014.

36. Steplewski A, Fertala J, Tomlinson R, Hoxha K, Han L, Thakar O, et al. The impact of cholesterol deposits on the fibrillar architecture of the Achilles tendon in a rabbit model of hypercholesterolemia. Journal of orthopaedic surgery and research. 2019;14(1):172.

37. van der Slot AJ, van Dura EA, de Wit EC, De Groot J, Huizinga TW, Bank RA, et al. Elevated formation of pyridinoline cross-links by profibrotic cytokines is associated with enhanced lysyl hydroxylase 2b levels. Biochim Biophys Acta. 2005;1741(1-2):95–102.

38. van der Slot AJ, Zuurmond AM, van den Bogaerdt AJ, Ulrich MM, Middelkoop E, Boers W, et al. Increased formation of pyridinoline cross-links due to higher telopeptide lysyl hydroxylase levels is a general fibrotic phenomenon. Matrix Biol. 2004;23(4):251–7.

39. Brickley-Parsons D, Glimcher MJ, Smith RJ, Albin R, Adams JP. Biochemical changes in the collagen of the palmar fascia in patients with Dupuytren’s disease. J Bone Joint Surg Am. 1981;63(5):787–97.

40. Paschalis EP, Verdelis K, Doty SB, Boskey AL, Mendelsohn R, Yamauchi M. Spectroscopic characterization of collagen cross-links in bone. J Bone Miner Res. 2001;16(10):1821–8.

41. Eyre DR, Paz MA, Gallop PM. Cross-linking in collagen and elastin. Annu Rev Biochem. 1984;53:717–48.

42. Rieppo L, Saarakkala S, Narhi T, Helminen HJ, Jurvelin JS, Rieppo J. Application of second derivative spectroscopy for increasing molecular specificity of Fourier transform infrared spectroscopic imaging of articular cartilage. Osteoarthritis Cartilage. 2012;20(5):451–9.

43. Gieroba B, Przekora A, Kalisz G, Kazimierczak P, Song CL, Wojcik M, et al. Collagen maturity and mineralization in mesenchymal stem cells cultured on the hydroxyapatite-based bone scaffold analyzed by ATR-FTIR spectroscopic imaging. Mater Sci Eng C Mater Biol Appl. 2021;119:111634.

44. Woessner JF. The determination of hydroxyproline in tisue and protein samples containing small proportions of this imino acid. ArchivBiochemBiophys. 1961;93:440–7.

45. Viele K, Berry S, Neuenschwander B, Amzal B, Chen F, Enas N, et al. Use of historical control data for assessing treatment effects in clinical trials. Pharmaceutical statistics. 2014;13(1):41–54.

46. Steplewski A, Fertala J, Cheng L, Wang ML, Rivlin M, Beredjiklian P, et al. Evaluating the Efficacy of a Thermoresponsive Hydrogel for Delivering Anti-Collagen Antibodies to Reduce Posttraumatic Scarring in Orthopedic Tissues. Gels. 2023;9(12):971.

47. Zhang J, Ding L, Zhao Y, Sun W, Chen B, Lin H, et al. Collagen-targeting vascular endothelial growth factor improves cardiac performance after myocardial infarction. Circulation. 2009;119(13):1776–84.

48. Liu J, Meng Z, Xu T, Kuerban K, Wang S, Zhang X, et al. A SIRPalphaFc Fusion Protein Conjugated With the Collagen-Binding Domain for Targeted Immunotherapy of Non-Small Cell Lung Cancer. Front Immunol. 2022;13:845217.

49. Tikhonov S, Babich O, Sukhikh S, Tikhonova N, Chernukha I. Synthesis of a new peptide and evaluation of its cytotoxicity and regenerative properties by prediction and in vitro experiment. Journal Of Applied Biology And Biotechnology. 2025;13(2):206–14.

50. Wei Z, Rolle MW, Camesano TA. Characterization of LL37 binding to collagen through peptide modification with a collagen-binding domain. ACS omega. 2023;8(38):35370–81.

