## Supplementary Materials for "Development of a Humanized Anti-Fibrotic Antibody Targeting Extracellular Collagen Assembly to Reduce Post-Traumatic Scarring"

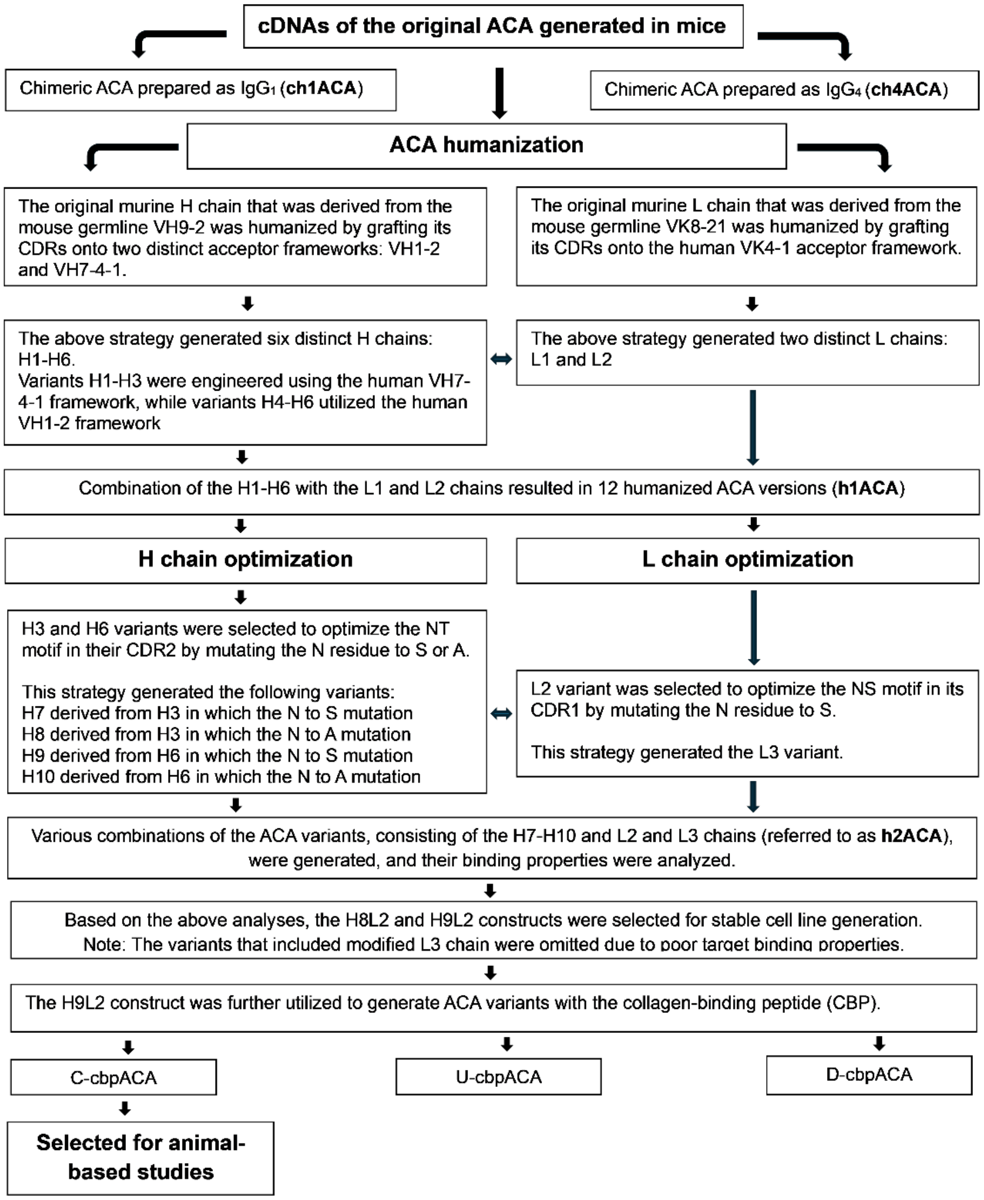


**Figure S1**. Flowchart summarizing the crucial steps in engineering humanized ACA variants.

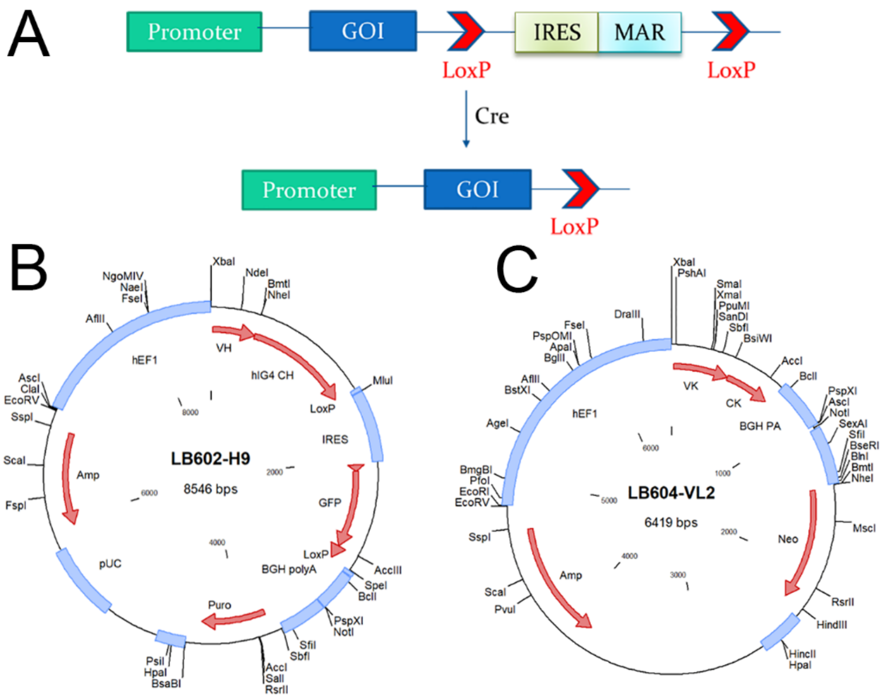


**Figure S2**. Schematic of the SwiMR expression system. (A) Bicistronic expression of a membrane-anchored reporter (MAR) is mediated by an internal ribosome entry site (IRES) downstream of the gene of interest (GOI). The IRES-MAR cassette is flanked by two LoxP sites for Cre recombinase recognition. Following Cre treatment, the sequence between the LoxP sites is excised from the chromosome. (B, C) Expression vector maps of the LB602-H9 and LB604-VL2 constructs, respectively.

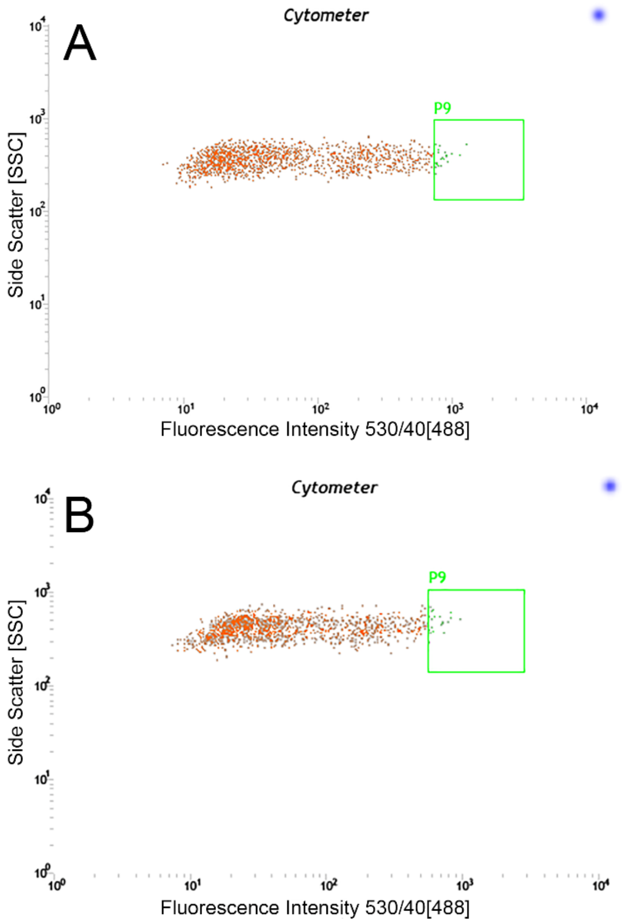


**Figure S3.** FACS isolation of high-producing cell lines. Stable pools of CHO-S cells, 44.42.2 and 44.42.3, were selected following transfection with LB602-H8/LB604-L2 and LB602-H9/LB604-L2 constructs, respectively. Cells were sorted based on GFP fluorescence (530/40 nm bandpass filter) versus side scatter (SSC). The top ~1% of cells exhibiting the highest GFP signal (gate P9) were isolated. (A) Flow cytometry plot for pool 44.42.2, with 1.43% of cells captured in the P9 gate. (B) Flow cytometry plot for pool 44.42.3, with 1.49% of cells captured in the P9 gate.

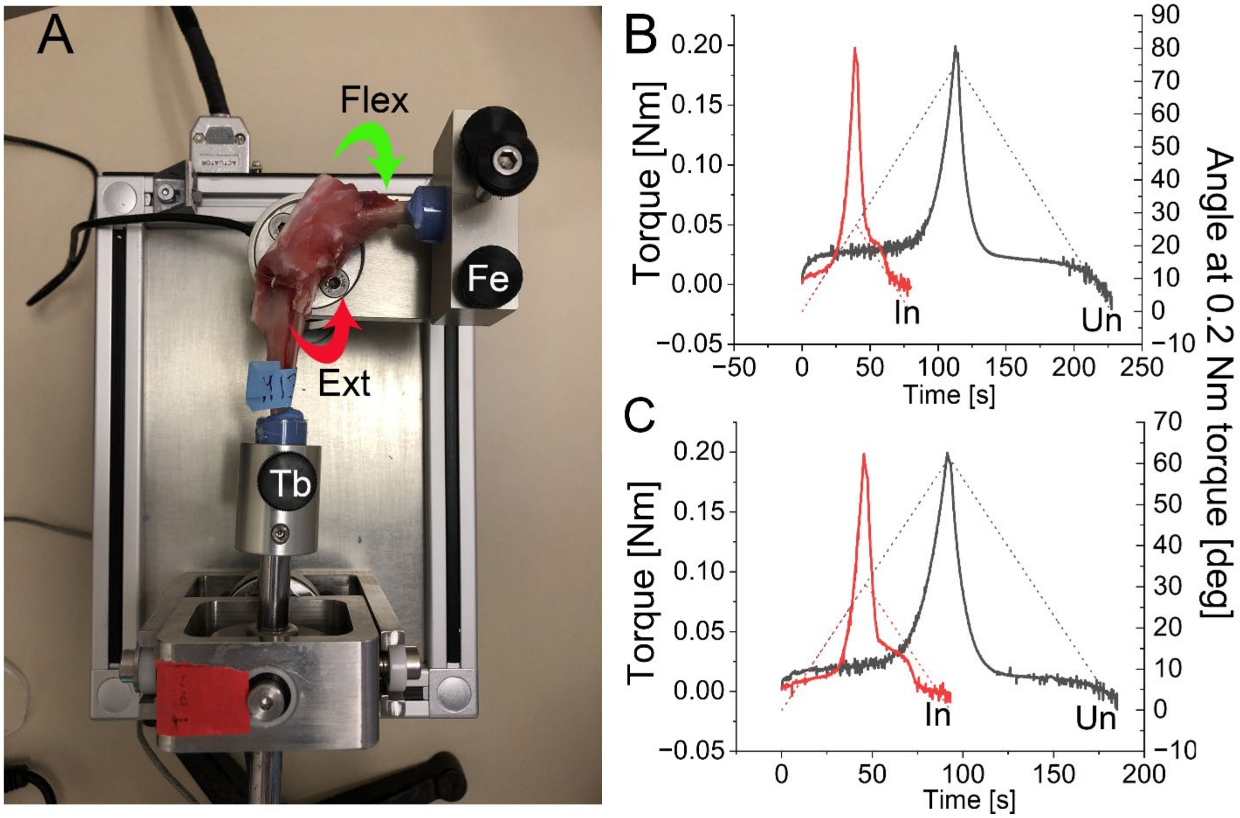


**Figure S4**. Measurement of knee flexion contracture in a rabbit joint model. (A) A custom-made biomechanical device utilized to measure joint stiffness. Tb, tibial clamp; Fe, femoral clamp. Arrows indicate the direction of rotation of the torque mechanism during the flexion (Flex) and extension (Ext) phases. (B, C) Representative joint stiffness measurements in two C-cbpACA-treated rabbits. Torque profiles of uninjured (Un, solid black curves) and injured (In, solid red curves) knees are shown during the extension (ascending) and flexion (descending) phases. Dotted curves indicate corresponding knee angles across varying torque levels. Flexion contracture was calculated using the knee angles of uninjured and corresponding injured limbs at a maximum applied torque of 0.2 Nm.

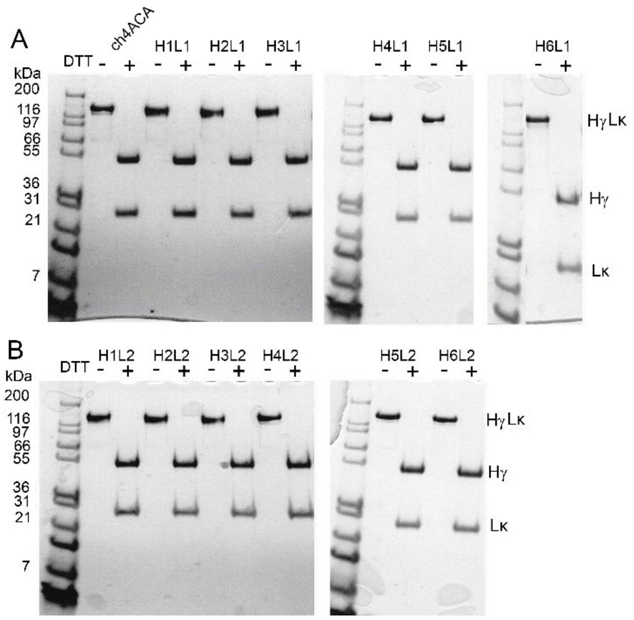


**Figure S5**. Electrophoretic analysis of first-generation humanized ACA variants. Antibodies comprised heavy chain variants (H1-H6) paired with light chain constructs (L1-L2). Samples were resolved by SDS-PAGE in the presence (+) or absence (-) of the reducing agent DTT. Migration patterns of the heavy (Hγ) and light (Lκ) chains are indicated. The chimeric ch4ACA antibody was included as a reference control.

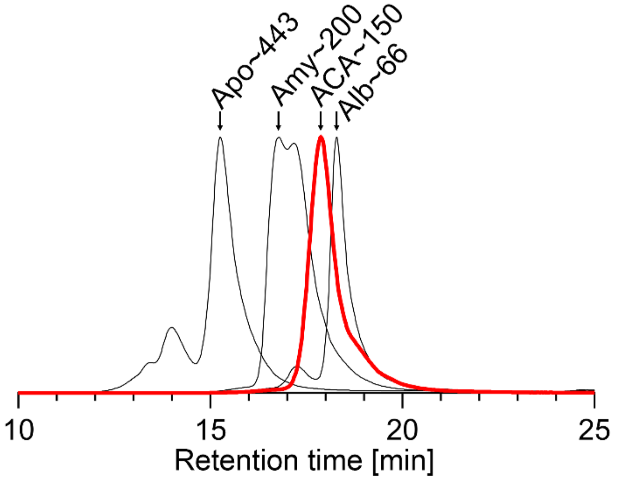


**Figure S6**. Representative size-exclusion chromatogram illustrating the expected elution profile of the ACA variants relative to molecular mass standards. Apo, aprotinin; Amy, amylase; Alb, albumin. Molecular masses of the indicated reference proteins are provided in kDa.

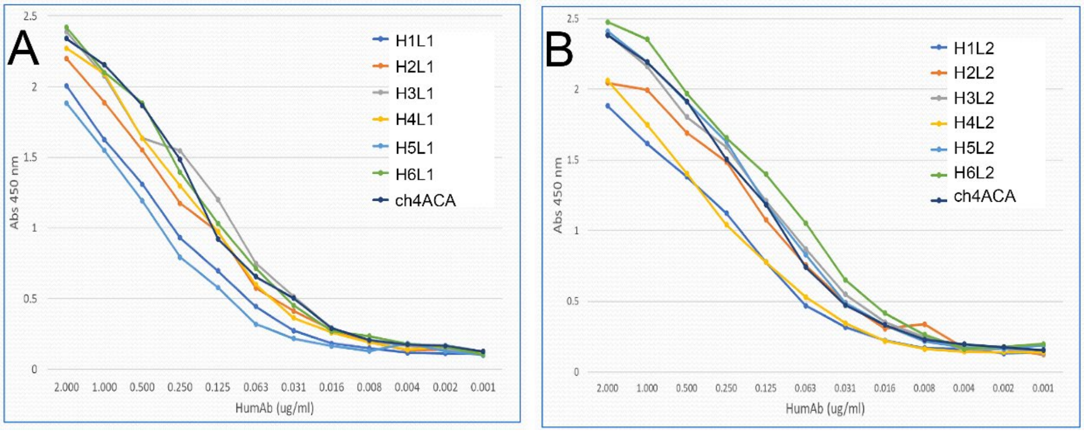


**Figure S7**. (A, B) Half-maximal effective concentration ($EC_{50}$) measurements for the binding of first-generation humanized ACA variants to the α2Ct target. Variants consist of heavy chains (H1-H6) paired with light chains (L1-L2). The chimeric ch4ACA construct was analyzed in parallel as a reference.

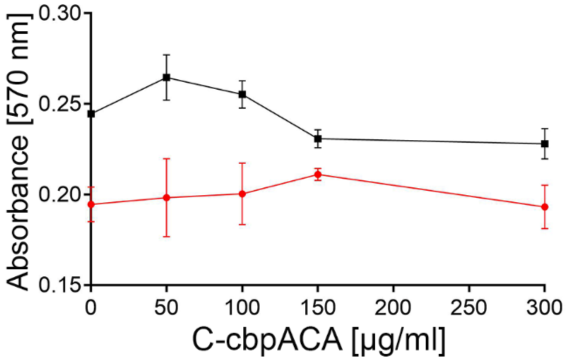


**Figure S8**. Viability of rabbit dermal fibroblasts (■) and human keloid-derived fibroblasts (●). Cells were cultured for 24 h in the presence of varying concentrations of C-cbpACA. Data represent the mean of 3 technical replicates (±SD).

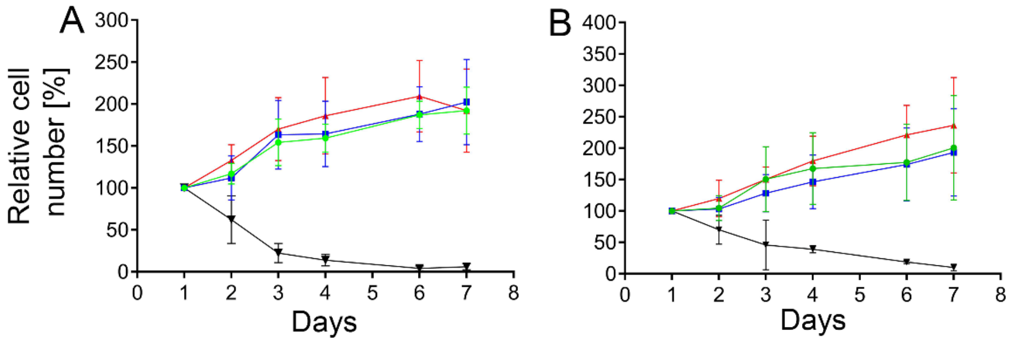


**Figure S9**. Proliferation assays of (A) rabbit dermal fibroblasts and (B) human keloid-derived fibroblasts cultured with 0, 100, or 200 μg/ml of mACA. Legend: 0 μg/ml (▲), 100 μg/ml (●), 200 μg/ml (■). Proliferation profiles of actinomycin D-treated cells are included as a negative control (▼). Data represent the mean of 3 technical replicates (±SD).

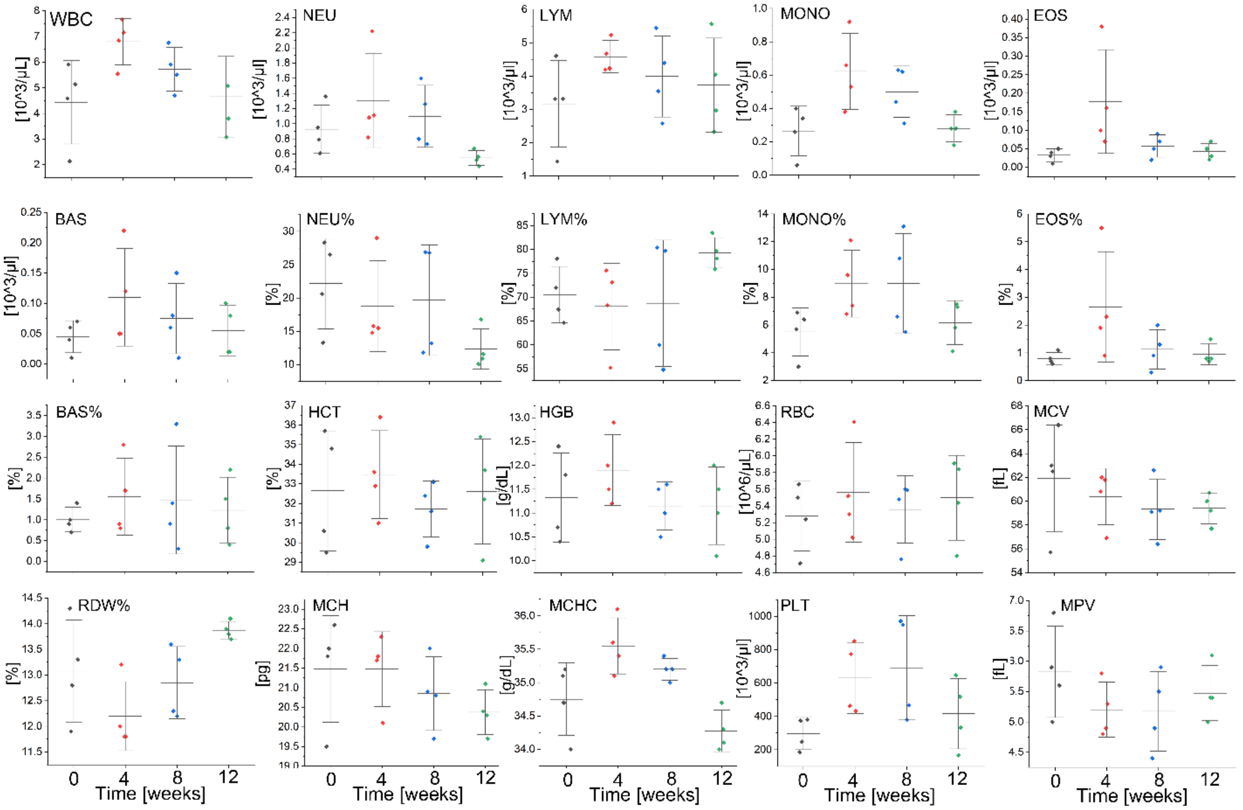


**Figure S10**. Complete blood count (CBC) parameters assessed prior to knee injury and at various post-operative time points. WBC; white blood cells, NEU; neutrophils, LYM; lymphocytes, MONO; monocytes, EOS; eosinophils, BAS; basophils, NEU %; percentage of neutrophils, LYM %; percentage of lymphocytes, MONO %; percentage of monocytes, EOS %; percentage of eosinophils, BAS %; percentage of basophils,

HCT; hematocrit, HGB; hemoglobin, RBC; red blood cells, MCV; mean corpuscular volume, RDW %; red blood cell distribution width, MCH; mean corpuscular hemoglobin, MCHC; mean corpuscular hemoglobin concentration, PLT; platelet count, MPV; mean platelet volume.

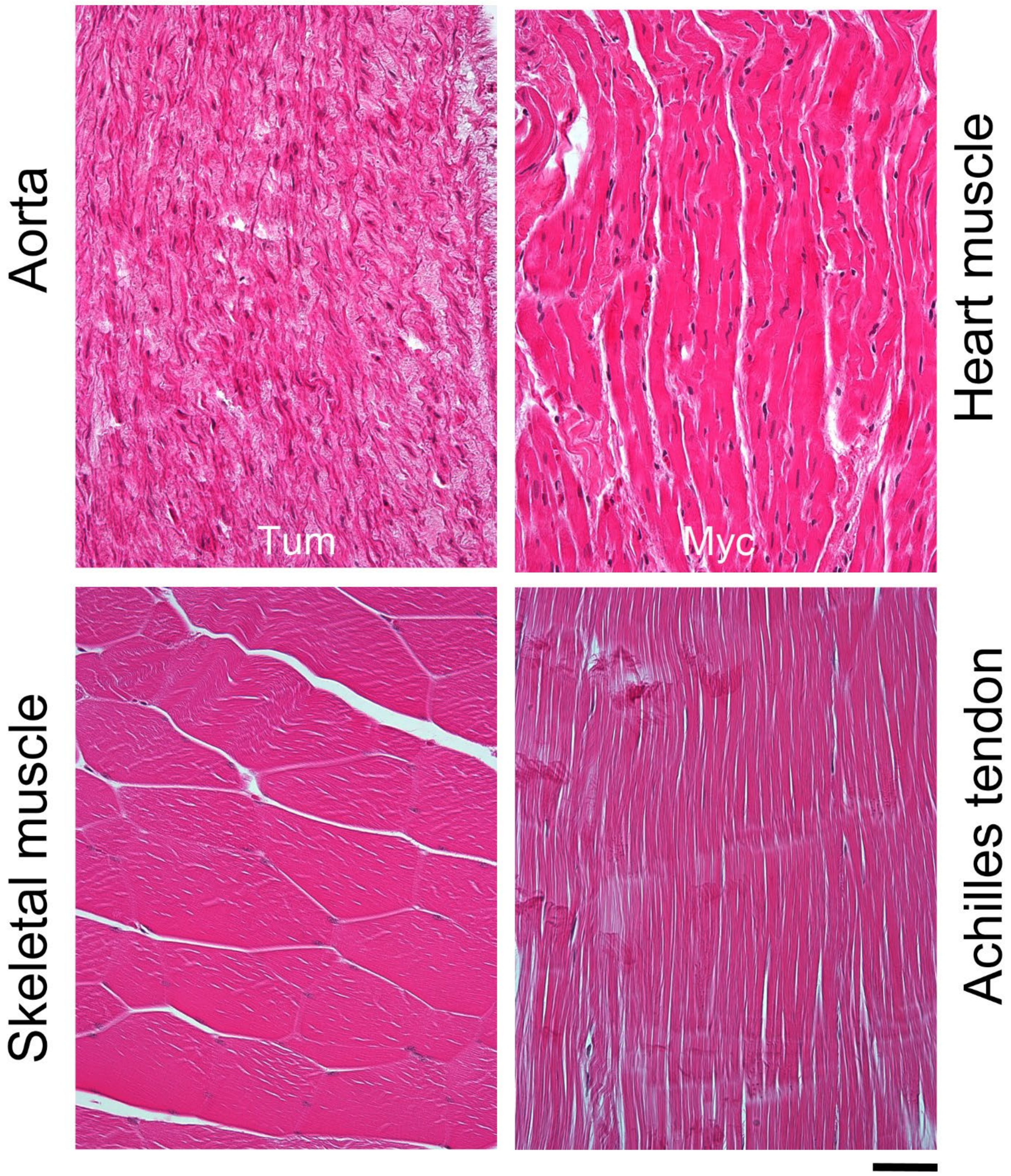


**Figure S11**. Histological evaluation of the aorta, heart, skeletal muscle, and Achilles tendon. Tum, tunica media; Myc, myocardium. Bar = 50 μm

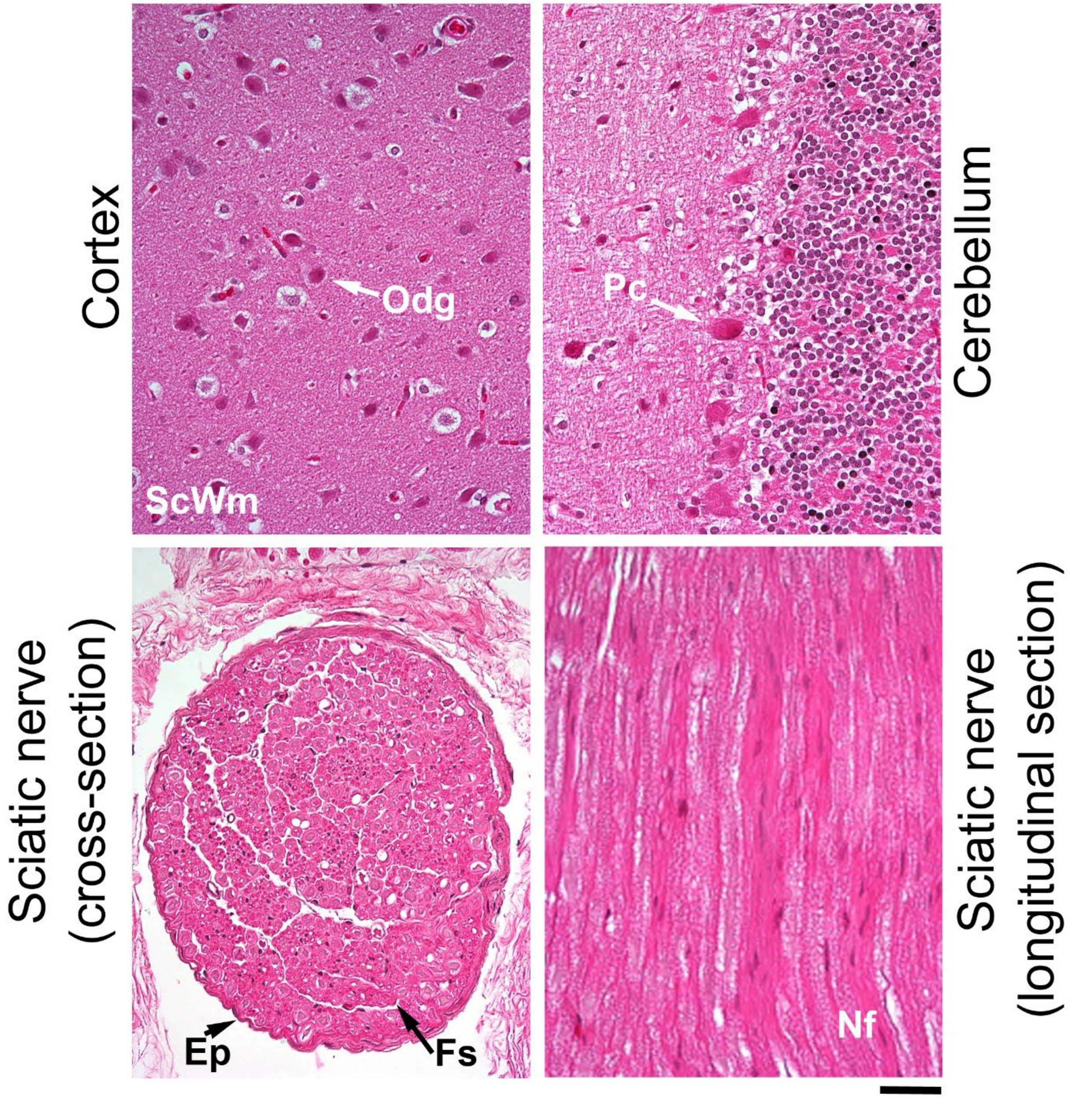


**Figure S12**. Histological evaluation of brain tissue and sciatic nerves. ScWm, subcortical white matter; Odg, oligodendroglia; Pc, Purkinje cells; Ep, epineurium; Fs, nerve fascicle. Bar = 50 μm

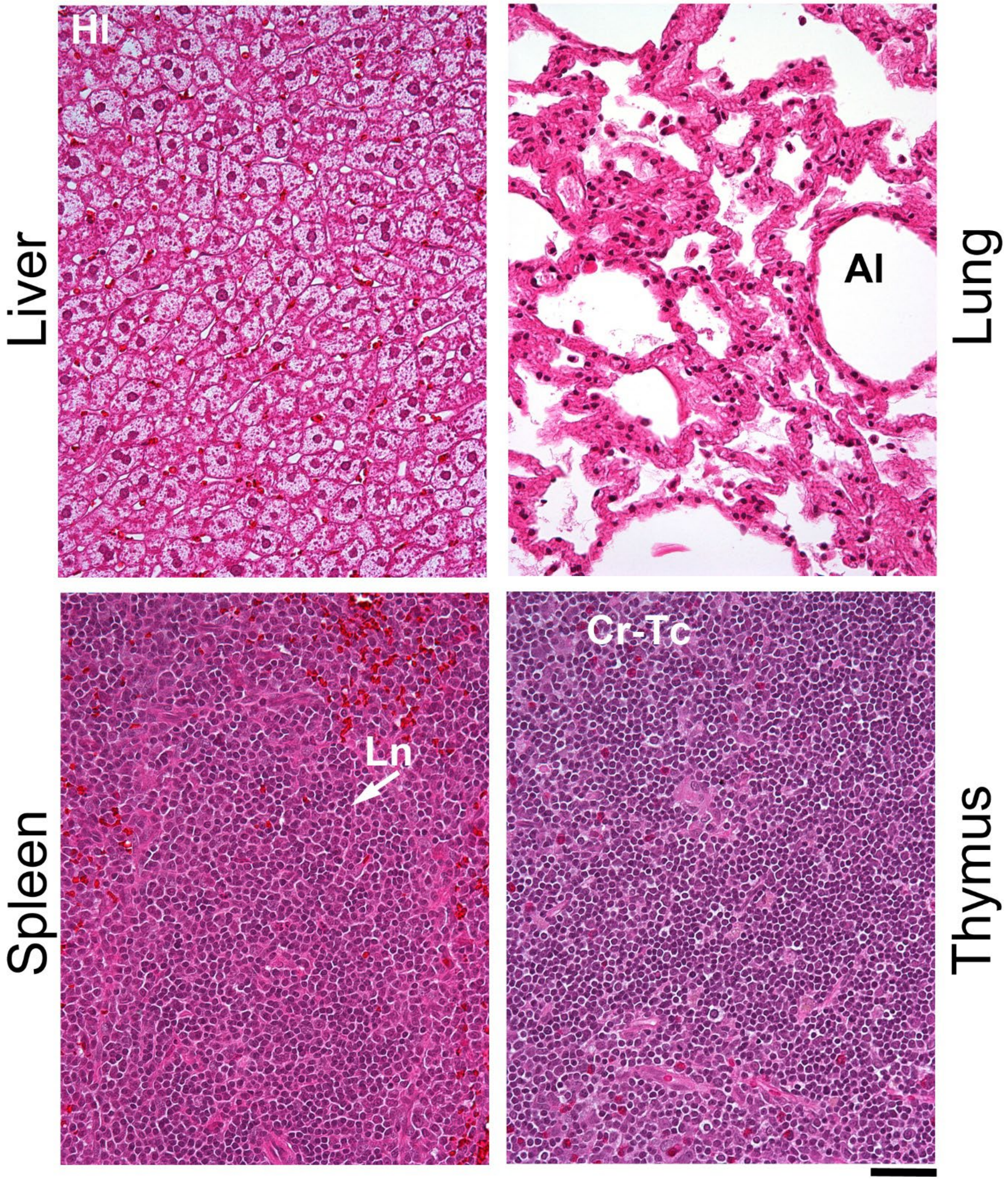


**Figure S13**. Histological evaluation of the liver, lung, spleen, and thymus. Hl, hepatic lobule; Al, alveolus; Ln, lymphoid nodule; Cr-Tc, cortical thymic cells. Bar = 50 μm

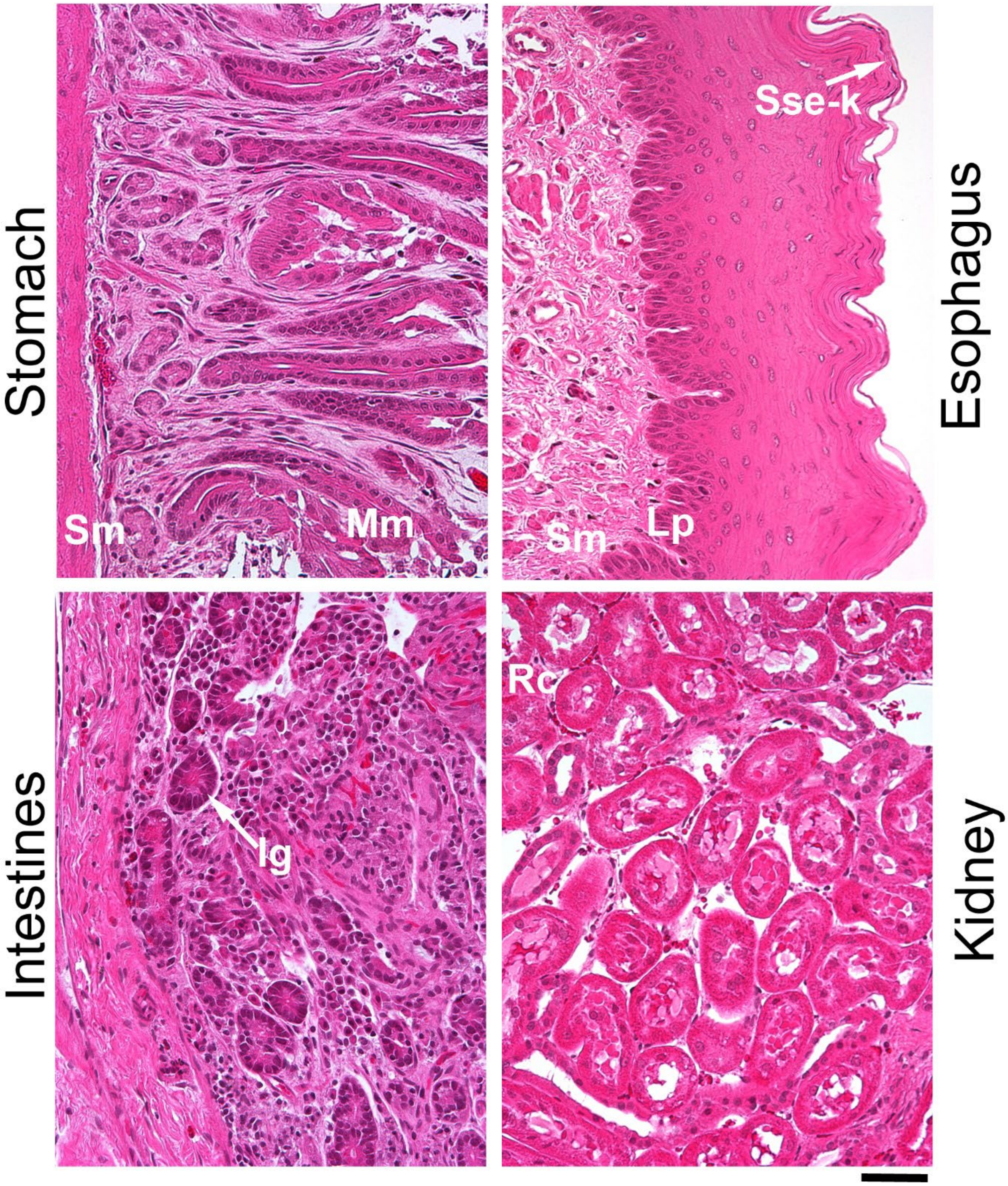


**Figure S14**. Histological evaluation of the stomach, esophagus, intestines, and kidney. Mm, mucous membrane; Sm, submucosa; Lp, lamina propria; Sse-k, stratified squamous epithelium-keratinized; Ig, intestinal glands; Rc, renal cortex. Bar = 50 µm

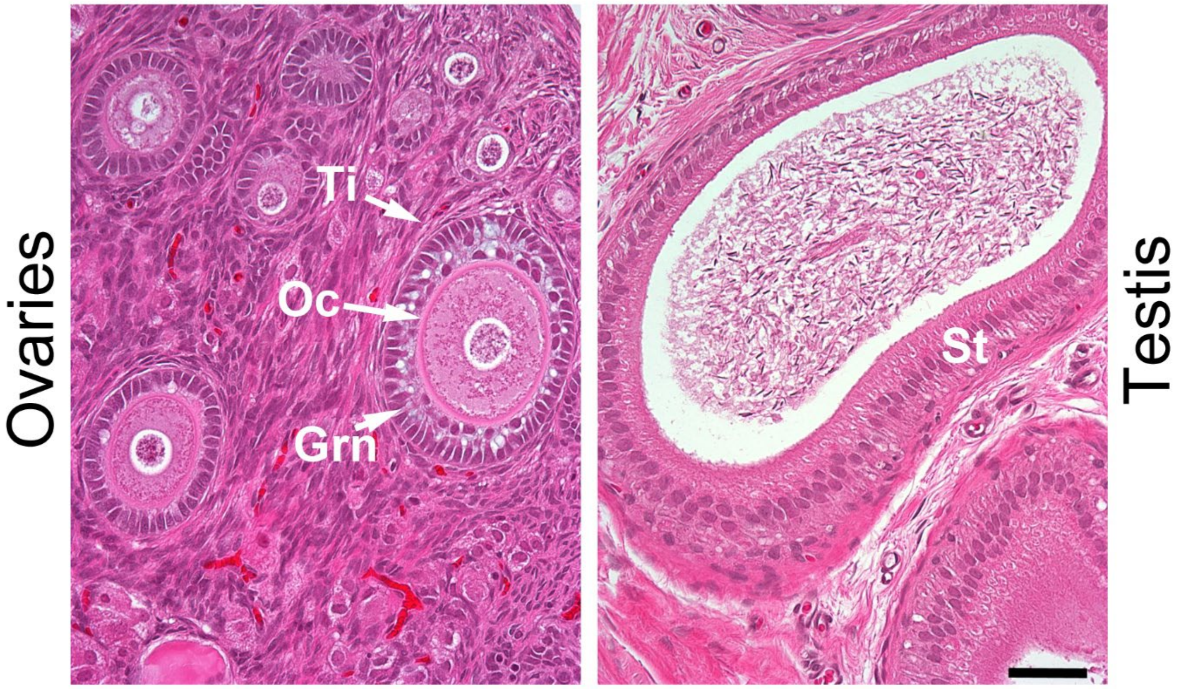


**Figure S15**. Histological evaluation of the ovaries and testes. Grn, granulosa cells; Ti, theca interna; Oc, oocyte; St, seminiferous tubule. Bar = 50 µm
